# The craniofacial shape of modern humans embodies genomic signatures of evolution, diversity, and clinical conditions

**DOI:** 10.64898/2025.12.29.696848

**Authors:** Seppe Goovaerts, Jay Devine, Nina Claessens, Sameer Gabbita, Jolien Deprest, Kaat Pauwels, Noah Herrick, Jaaved Mohammed, Aurélien Mounier, Tamar Sofer, Hannah K. Long, Ullrich Bartsch, Benedikt Hallgrímsson, Sarah J. Lewis, Stephen Richmond, Sarah Bauermeister, Susan Walsh, John R. Shaffer, Mark D. Shriver, Sahin Naqvi, Joanna Wysocka, Toomas Kivisild, Seth M. Weinberg, Peter Claes

## Abstract

Human craniofacial morphology is a hallmark of our species’ diversity and evolutionary history, shaped by adaptation, introgression, and global dispersal. Cranial globularization and chin emergence are well-documented morphological transformations whose genetic basis remains poorly understood, whereas Neandertal introgression is primarily documented through genomic evidence. How these evolutionary phenomena relate to craniofacial variation in present-day humans remains largely unresolved. Here, we leverage 3D craniofacial data from over 50,000 UK Biobank participants and employ a multivariate, multiscale genome-wide association approach to define axes of variation aligned with inter-population allele frequency shifts, evolutionary processes, and clinical conditions. We identify continuous craniofacial trends within our cohort that mirror global patterns of genetic diversity, indicating that facial differences between human populations arise at the phenotypic axes already present within a single population. We further demonstrate that modern human-derived alleles underlie the origins of the human chin by reducing midfacial projection relative to other hominins and reveal the persistent effects of Neandertal introgression on craniofacial diversity today. We also model genetically informed endophenotypes for orofacial clefts, obstructive sleep apnoea, and myopia. These findings provide insights into our species’ evolutionary history and endophenotypes of clinical conditions and establish a framework for contextualizing craniofacial diversity into biologically meaningful axes of variation relevant to diverse scientific disciplines.

## Main

Human craniofacial morphology is a hallmark of our species’ diversity and evolutionary history. Its complexity arises from the coordinated development of multiple tissues^1,2^ and is governed by a highly polygenic genetic architecture^3^. The emergence of novel alleles during hominin evolution allowed natural selection and neutral processes to drive coordinated shifts in allele frequencies that distinguish *Homo sapiens* from other hominins. On the morphological level, this occurred most conspicuously via globularization of the braincase^4^ (with concomitant changes in the cranial base and face^5^), in parallel to the derived emergence of a triangular mental eminence (i.e., the modern human chin), a feature whose adaptive significance remains a topic of debate^6^. On the genomic level, gene-regulatory divergence at human-accelerated regulatory elements in cranial neural crest cells (CNCCs), a transient multipotent population of cells that contribute to the majority of craniofacial structures^1^, has pointed to regulatory evolution as a major driver of craniofacial change^7,8^. Past human migrations have further shaped craniofacial variation by creating allele frequency gradients via long-term genetic drift, serial founder events^9^, and local adaptive pressures^10^. These gradients produced phenotypic contrasts that have been misinterpreted as biological evidence for socially constructed “race” categories, despite the underlying genetic and phenotypic continuity across populations^11^.

Indeed, both derived and ancestral alleles persist in modern populations as single nucleotide polymorphisms (SNPs) that can influence craniofacial shape^12–21^. Some of these SNPs may additionally alter the genetic liability for congenital anomalies, such as orofacial clefts^22^, or contribute to complex health conditions in which craniofacial shape is a risk factor^23^. Another source of genetic variation affecting craniofacial shape^16,24^, as well as other complex traits^25^, stems from introgressed DNA segments that persist in most contemporary populations outside Africa since interbreeding the with Neandertals ∼50 *kya*^26^.

Advances in our understanding of craniofacial variation related to evolutionary processes^27–30^ and complex health conditions^23,31^ have largely stemmed from studies focusing on either morphology or genetic variation. However, these efforts have rarely integrated directional genomic signals with craniofacial shape, leaving a key biological question unresolved^32^: how do evolutionary, demographic, and health-related genomic signals shape morphological diversity in present-day humans? Here, using over 50,000 whole-head magnetic resonance (MR) images in the UK Biobank (UKBB), we create a large dataset of linked genetic and phenotypic data and address two major gaps: (i) expanding the catalogue of genetic variants associated with human craniofacial morphological variation through a multivariate, multiscale genome-wide association study (GWAS), and (ii) introducing a context-aware framework encompassing evolutionary and demographic processes and genetic susceptibility to clinical conditions.

## Results

### An expanded catalogue of genetic variants associated with craniofacial shape

We extracted craniofacial surfaces from high-resolution MR images (*n* = 50,622) in the UKBB (**Fig 1a**, Methods), using unrelated individuals of self-reported White-British ethnicity with similar genetic ancestry and a median age of 65. To capture shape variation, we mapped dense, homologous quasi-landmarks (*n* = 14,903) onto each individual, then decomposed these configurations into 67 hierarchical segments, enabling global and local shape analyses (**Fig 1a**; see Fig S1 and Supplementary Note for segment numbers and definitions). The configurations of each segment were Procrustes superimposed and summarized into principal components (PCs; *n* = 13–145) explaining 98% of variance after adjustment of shape data for covariates (sex, age, age-squared, anthropometric measures, scanner parameters, assessment centre/date, size, and the first ten genomic PCs). Per segment, we conducted a multivariate GWAS to test for association between its full set of PCs and each of the 8,922,008 SNPs with a minor allele frequency (MAF) greater than 1%, spanning the autosomes and X-chromosome.

**Fig 1.**
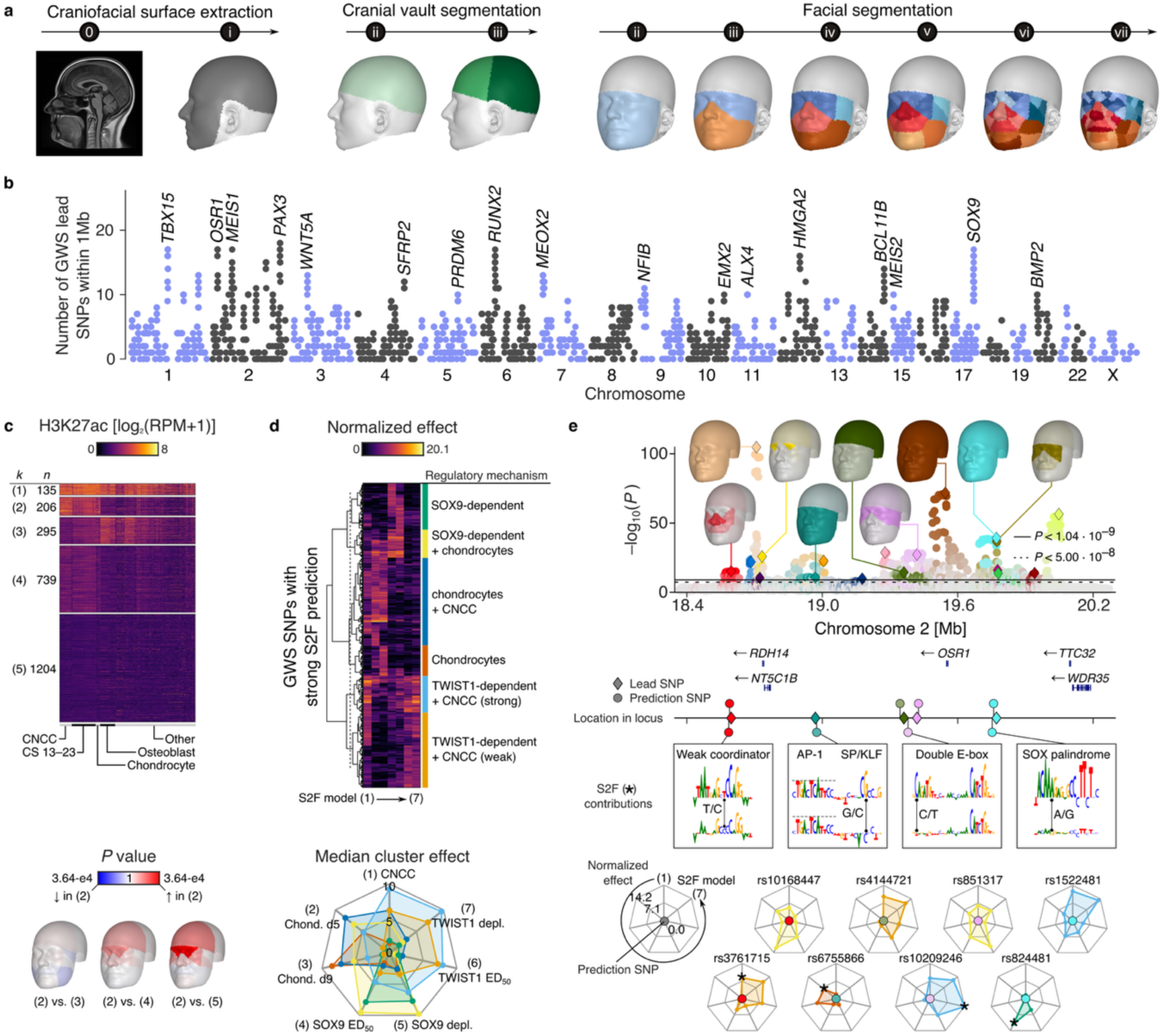
Genome-wide discovery of global-to-local effects on craniofacial shape. **(a)** Extraction of the craniofacial surface from magnetic resonance (MR) images and segmentation into local segments. Hierarchical levels are indicated with roman numerals; the full surface is indicated “i”. **(b)** Brisbane plot showing the number of conditionally independent genome-wide significant (GWS) lead SNPs within 1Mb of each GWS lead SNP (*n* = 2579), i.e., lead SNP density. Non-significant and non-lead SNPs are not shown. **(c)** H3K27ac signal in the 10 kb bin surrounding each GWS lead SNP. SNPs are grouped into five clusters based on activity across cell and tissue types. For SNP-cluster pairs with significant differences in effect localization (segment-R^2^; permutation MANOVA with post-hoc testing), segment-wise comparisons were performed using the Mann-Whitney *U* test. *P* values are shown on a log scale. **(d)** Clustering of functional predictions (*P* < 0.01) of GWS SNPs (*n* = 1635) located near cranial neural crest cell (CNCC) ATAC peaks (≤750 bp), obtained using deep learning sequence-to-function (S2F) prediction models based on ChromBPNet. Columns represent seven S2F models predicting: (1) steady-state ATAC signal in CNCCs, (2) chondrocytes at day 5 (d5), (3) chondrocytes at day 9 (d9); and SOX9- (4, 5) and TWIST1-dependent (6, 7) effects on ATAC signal in CNCCs, characterized by (4, 6) median effective dosage (ED50) and (5, 7) full SOX9/TWIST1 depletion. Radar plot shows median effects per cluster. **(e)** LocusZoom plot of a ∼1.8 Mb region surrounding *OSR1* containing 18 independent GWS lead SNPs (diamonds). Phenotypic effect localization is indicated by colour matching the lead SNP, with intensity reflecting –log10(*P*) across craniofacial regions. For non-lead SNPs, colour indicates the most strongly linked lead SNP and intensity reflects LD. Radar plots show S2F predictions for selected SNPs in genomic order, with colour indicating the regulatory cluster in (d); centre sphere colour matches SNP colour in the LocusZoom plot. Contribution plots indicate how these SNPs influence S2F scores by creating transcription factor binding motifs. The model is indicated by ‘*’.

Across the 67 segments, we identified between 37 and 2054 conditionally independent genome-wide significant (GWS; *P* < 5e-8) lead SNPs which were aggregated into a total of 2579 conditionally independent GWS lead SNPs located at 1175 non-overlapping genomic regions (**Fig 1b**; see Fig S2a for a Manhattan plot and Supplementary Data 1 for the full list of GWS lead SNPs). Of these, 2108 were also study-wide significant at an adjusted ⍺ level of 1.04e-9 which considers the effective number of tests per SNP (Methods). Using three independent GWAS datasets, including whole-face meta-analyses by White *et al.* (*n* = 8246) and Zhang *et al.* (*n* = 9674), as well as the cranial vault GWAS by Goovaerts *et al.* (*n* = 4198), we replicated 973 (38.6%), 728 (30.0%), and 408 (16.1%) of all GWS lead SNPs, respectively, at *P* < 0.05 (Methods; Supplementary Data 2). Across the same datasets, 652 (25.9%), 425 (17.5%), and 122 (4.8%), GWS lead SNPs respectively were also significant at a 5% false discovery rate (FDR). Non-replicating SNPs still showed a strong enrichment of signal in both facial datasets (Fig S2b), highlighting the substantial power increase in this work over previous studies. Linkage disequilibrium score regression^33^ (LDSC) intercepts ranged between 0.98 and 1.04 (mean = 1.01; s.d. = 0.01) confirming the absence of residual confounding. The LDSC common-variant heritability (h^2^) of craniofacial shape was estimated as 22.4%, with a similar h^2^ for the whole face (23.4%) and whole cranial vault (22.1%), and up to 27.5% for the nose (facial segment 8; see Supplementary Data 3 for other segments).

A high density of GWS SNPs was observed near genes encoding key craniofacial transcription factors (TFs) including *TBX15*, *OSR1*, *MEIS1*, *RUNX2*, and *SOX9*, where GWS lead SNPs shared their location with up to 17 other conditionally independent GWS lead SNPs (**Fig 1b**). The highest density was observed in the region surrounding *PAX3* with 19 independent association signals. Overall, 2333 (90.5%) GWS lead SNPs had at least one secondary signal within 1 Mb, while the average was 4.59, often with distinct association patterns across anatomical regions or segmentation levels (Supplementary Data 1).

Re-analysis of chromatin immunoprecipitation followed by sequencing (ChIP-seq) data for the K27ac mark on histone 3 (H3K27ac) was used to investigate cell type-specific regulatory activity in the vicinity (10 kb) of the 2579 GWS lead SNPs. We observed an enrichment of H3K27ac signal in cranial neural crest cells (CNCCs; *P* = 2.06e-6; *n* = 12), embryonic craniofacial tissue from Carnegie stages 13–23 (CSs; *P* = 7.52e-13; *n* = 22), osteoblasts (*P* = 1.13e-4; *n* = 15), and CNCC-derived chondrocytes (*P* = 4.75e-3; *n* = 4) relative to other, less related cell types (*n* = 74; right-tailed Mann-Whitney *U* test). Clustering the SNPs based on their H3K27ac signal revealed a group of SNPs (*n* = 206; group 2 in **Fig 1c**) with high activity in CNCCs, CSs, and CNCC-derived chondrocytes, while having low activity in osteoblasts and other cell types (**Fig 1c**). Meanwhile, the opposite was observed for group three (*n* = 295), which showed high activity in osteoblasts and low activity in other cell types, except for fibroblasts (in “other”). Furthermore, individual SNPs across the five groups varied in the patterns and amounts of phenotypic variance explained across the 67 craniofacial segments (*P* = 1.66e-2; *F* test using residual randomization^34^). Pairwise differences were significant (*F* test using residual randomization and Holm adjustment) between groups two and three (*Padj* = 7.6e-3), two and four (*Padj* = 1.0e-3), and two and five (*Padj* = 1.1e-3). Relative to groups four and five, the SNPs in group two explained more shape variance around regions tied to the frontonasal prominence, including the nose, orbits, and frontal bone (**Fig 1c**; minimum *P* = 3.64e-4; two-tailed Mann-Whitney *U* test), whereas SNPs in group three explained suggestively more variance in the mandible relative to those in group two (minimum *P* = 2.18e-2). These results demonstrate that cell type-specific activity in the direct vicinity of our GWS lead SNPs can be linked to craniofacial region-specific effects.

To investigate *cis*-regulatory mechanisms underlying our GWAS loci in greater detail, we utilized seven sequence-to-function (S2F) deep learning models introduced previously^35^ and based on the ChromBPNet^36^ backbone to predict the functional impact of allele substitutions at GWS SNPs (*n* = 272,953). Three models predicted fold-changes in steady-state chromatin accessibility, as measured by ATAC-seq in CNCCs and CNCC-derived chondrocytes at differentiation day 5 (d5) and day 9 (d9). Four additional models predicted how concentrations of dosage-sensitive TFs, SOX9 and TWIST1, influence chromatin accessibility in CNCCs. Specifically, we modelled (i) fold-change in accessibility upon full TF depletion, where large effects suggest direct TF targets, and (ii) changes in median effective dosage (ED50), i.e., the TF concentration at which accessibility is halved, where lower ED50 indicates dosage sensitivity and higher ED50 reflects buffering.

Clustering SNPs with strong predictions (*P* < 0.01 in ≥ 1 S2F model, *n* = 1635) revealed distinct *cis*-regulatory mechanisms (**Fig 1d**). For example, some SNPs were predicted to affect steady-state accessibility in CNCCs, chondrocytes, or both, without altering TF dosage sensitivity, whereas others were predicted to primarily modulate SOX9 responsiveness. Strong predictions for TWIST1 dosage sensitivity often coincided with strong predictions for CNCC accessibility, and one cluster of SNPs was predicted to modulate both SOX9-dependency and chondrocyte accessibility, consistent with known TF functions^37,38^. Notably, we observed that many GWAS loci harbour SNPs from multiple clusters. For example, a ∼1.8 Mb region near *OSR1* harboured SNPs from four out of the six major clusters, most in strong LD with a nearby lead SNP (**Fig 1e**). For a subset of these SNPs, we derived base-resolution contribution scores for the immediately surrounding sequence and assessed how each allele influenced these sequence contributions (Fig 1e). For example, rs3761715 alters a weak ‘Coordinator’ motif bound by TWIST1, which is predicted to affect steady state ATAC-seq signal in CNCCs. In contrast, rs10209246 alters a double E-box motif, previously shown to be bound by TWIST1^39^ and especially predictive of sensitivity to TWIST1 dosage^35^. rs824481 alters a SOX9 palindrome motif, identified previously to be highly predictive of SOX9 responsiveness^35^. rs6755866, which is predicted to alter steady-state ATAC-seq signal in chondrocytes, alters a GC-rich sequences typically bound by the SP/KLF TFs. Notably, this alteration also reduces the contribution score of a nearby AP-1 binding site, a TF known to regulate chondrogenesis^40^, suggesting that this SNP modifies the larger context around the AP-1 site. Consistent with prior observations, the SNPs near the centre of ATAC-seq peaks were predicted to primarily affect steady-state accessibility, whereas those at the flanks modulate transcription factor responsiveness^35^ (Fig S1c). At loci containing independent GWS lead SNPs with distinct phenotypic effects, exemplified by the OSR1 locus, the associated variants are predicted to alter *cis*-regulatory activity in a cell type- and TF dosage-dependent manner (Fig 1e). This suggests a complex spatiotemporal pattern of *cis*-regulation at key craniofacial genes, facilitating pleiotropic roles of these genes in craniofacial morphology.

### Intra-cohort trait continuity extends to global diversity

Geographical structuring of human populations, prompted by past migrations, has led to shifts in allele frequencies driven by a combination of neutral processes and local adaptive pressures^9^. By using the 1000 Genomes Project^41^ (1KG) dataset to identify frequency-shifted SNPs at craniofacial shape loci and subsequently modelling their joint effect, we assessed whether phenotypic continuity within our cohort extends to encompass global patterns of craniofacial diversity.

For 24 1KG populations, spanning five continents, a set of quasi-independent (r^2^ < 0.1) craniofacial predictor SNPs (*n* = 30–1848; Supplementary Data 5) was selected based on our GWAS signal and allele frequency differences relative to the British from England and Scotland (GBR; Fisher’s exact test; right tailed χ^2^). To model the effects of each set of frequency-shifted SNPs on craniofacial morphology, we first fit a multivariate linear model on the genotypes and phenotypes of our UKBB sample (*n* = 50,622; Methods). We then evaluated this model at population-average allele counts, producing two opposing point predictions in the phenotypic principal component analysis (PCA) space, uniquely defining a context-aware axis of shape variation associated with the directionality of the population divergence. The population-aligned axes captured 6.72–14.14% of shape variance (Supplementary Data 5), significantly exceeding random axes (mean = 0.69%; s.d. = 0.18%; *P* < 1e-6; one-tailed empirical test). Variation along these axes exhibited approximately normal distributions (**Fig 2a**), suggesting that they represent broad phenotypic trends rather than outlier-driven effects.

**Fig 2.**
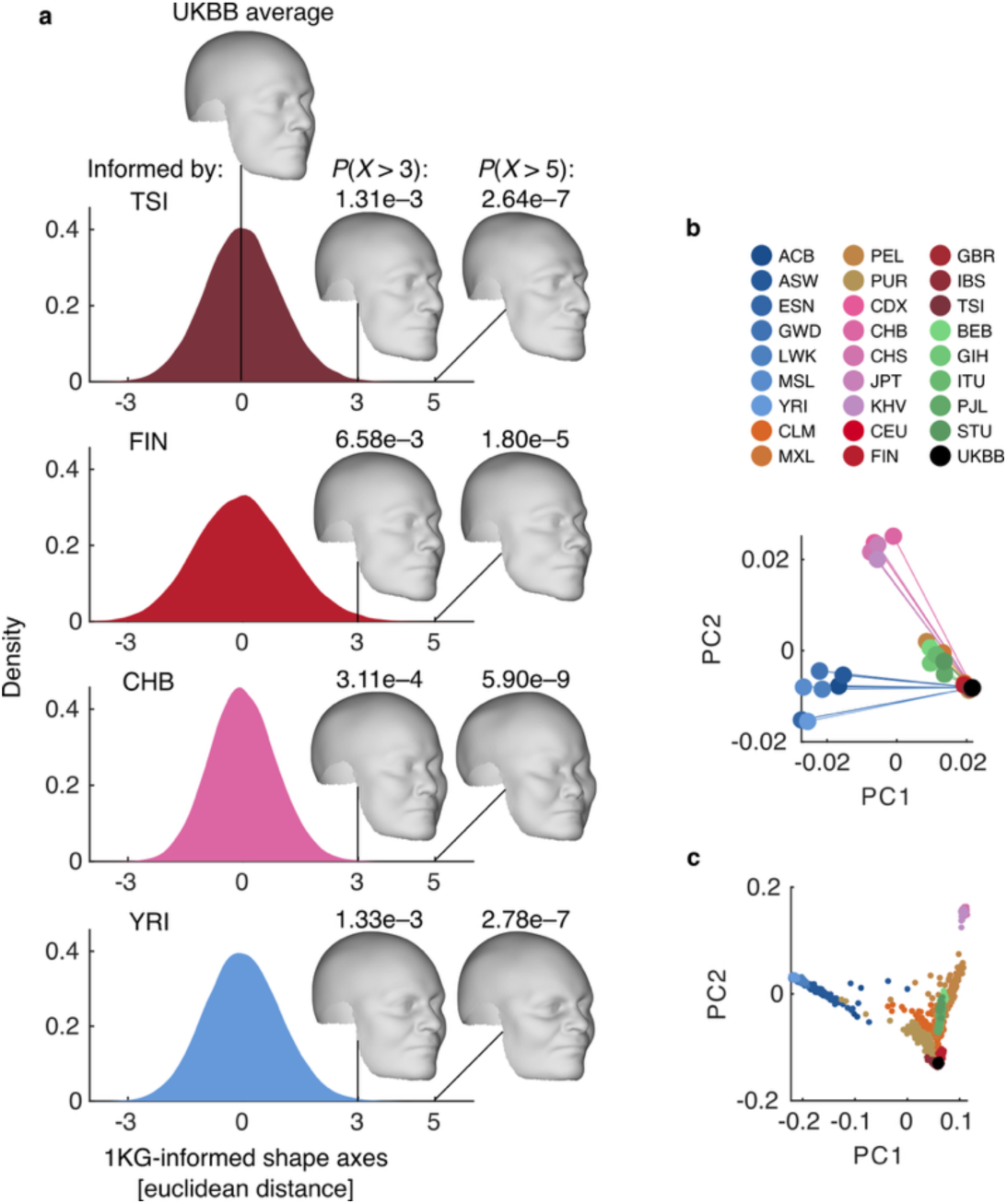
Alleles in UKBB GWAS sample capture global patterns of craniofacial diversity. **(a)** Shape axes in the UKBB GWAS sample informed by 1000 Genomes Project (1KG) labels [Toscani in Italy (TSI); Finnish in Finland (FIN); Han Chinese in Beijing, China (CHB); and Yoruba in Ibadan, Nigeria (YRI)]. Distributions show the one-dimensional projections of UKBB individuals onto the shape axes. Shape visualizations are made at three and five units from the mean shape to illustrate expected variation at the distribution margins and beyond, respectively. Probabilities indicate the likelihood that a shape’s projection extends at least this far along the axis. **(b)** Phenotypic subspace formed by 1KG-informed phenotypes (*n* = 24) and the UKBB mean shape. **(c)** Genetic ancestry space of 26 1KG populations (*n* = 2504) alongside randomly sampled UKBB individuals (*n* = 2500).

Expected shapes at the distribution margins exhibited moderate phenotypic differences relative to the UKBB mean shape as exemplified using the axes aligned to frequency shifts between GBR and Toscani in Italy (TSI), Finnish in Finland (FIN), Han Chinese in Beijing, China (CHB), and Yoruba in Ibadan, Nigeria (YRI) (**Fig 2a**). Moving further along these axes revealed phenotypes whose probability of occurring became exceedingly low under the multinomial allele frequency distribution of our UKBB sample (**Fig 2a**). For example, the CHB-aligned axis captured a less prominent nasal bridge with increased midfacial breadth and relatively pronounced zygomatic regions (**Fig 2a**). The YRI-aligned axis captured a wider nasal alae and a broader alar base (**Fig 2a**), features previously associated with variation in nasal morphology across global populations and linked to climate adaptation in prior studies^10^. Additionally, the midface was slightly more prognathic, and the cranial vault displayed a more brachycephalic configuration (**Fig 2a**). The TSI-aligned axis was characterized by a more convex nasal profile with a prominent bridge and a downward-oriented tip, along with a narrower facial width, a pronounced mental region, and a more inclined frontal bone (**Fig 2a**). Finally, the FIN-aligned axis exhibited a relatively square mandibular contour, elevated zygomatic arches, and a lower nasal bridge (**Fig 2a**).

When projecting the 1KG-aligned axes (*n* = 24) into a phenotypic subspace using PCA (Methods), the first two PCs formed a triangular configuration that closely mirrored the structure of the first two genetic PCs (**Fig 2b–c**). This correspondence indicates that 1KG-aligned phenotypic axes capture global patterns of craniofacial variation present in our UKBB cohort. Moreover, populations that were closer aligned geographically and genetically were projected onto more closely aligned phenotypic axes, despite SNP selection and model fitting being performed independently for each population. This concordance demonstrates robustness and validity of the axes. Collectively, these findings show that 1KG-aligned axes capture meaningful craniofacial variation within our UKBB European ancestry cohort, aligned with gradients of population divergence, revealing patterns that trace back to shared ancestry and historical dispersal.

### Derived alleles underlie the emergence of a modern human chin

We next sought to investigate the evolutionary origins of a triangular mental eminence, a feature that emerged in *Homo sapiens* and that is absent in other hominins. In particular, we tested whether derived alleles collectively explain chin prominence and how these alleles may have simultaneously influenced other integrated anatomical structures. For this, we obtained the most likely ancestral and derived states of facial GWAS SNPs (whole face; *P* < 5e-8), previously inferred based on a six-way primate multisequence alignments^42^ (**Fig 3a**). Much like the axes capturing population divergence, we applied our context-aware approach to define an ancestral-derived shape axis based on the joint effect of homozygous ancestral and derived states across 1958 quasi-independent (r^2^ < 0.1) SNPs in our UKBB sample (*n* = 50,622; Methods). The derived end of this axis revealed a prominent chin accompanied by a flatter and retrognathic midface (**Fig 3b**), while at the ancestral end, the chin had almost disappeared, and the face was more prognathic. These results confirm that the human chin can be explained by derived alleles and suggest that the same alleles contribute to reduced midfacial projection compared to other hominins.

**Fig 3.**
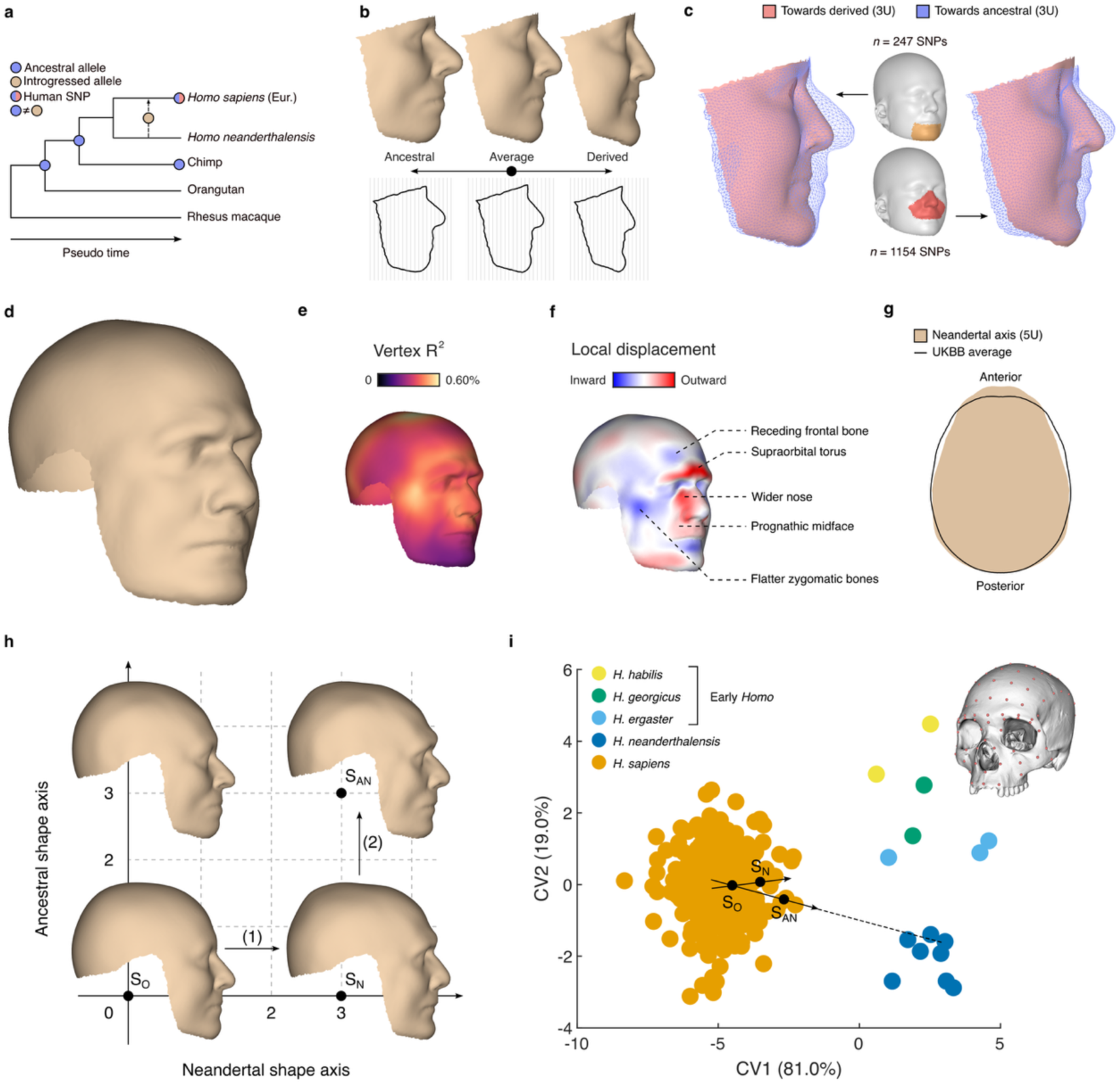
Genomic signatures of evolution and introgression reflected in present-day craniofacial diversity. **(a)** Conceptual schematic of hominin lineages relative to a macaque outgroup, illustrating “ancestral,” “derived,” and “Neandertal introgressed” allele categories. **(b)** Axis of facial variation associated with ancestral or derived states at craniofacial GWAS loci (*n* = 1958 SNPs). Facial renderings and outlines depict expected shapes at the extremes of the UK Biobank distribution (±3 Euclidean distance units). **(c)** Renderings compare facial phenotypes associated with ancestral and derived states modelled from SNPs significant for chin (*n* = 247) or midfacial (*n* = 1154) morphology. **(d)** Expected shape at the UKBB distribution margins (±3 Euclidean distance units) along the Neandertal-aligned shape axis modelled from introgressed SNPs (*n* = 79). **(e)** Vertex-wise map of variance explained by introgressed SNPs. **(f)** Heatmap showing local displacements of the shape in (d) relative to the UK Biobank mean; annotated with known anatomical features. **(g)** Cranial vault shape effects captured by the Neandertal-aligned shape axis magnified beyond the UKBB distribution margins (5 Euclidean distance units). **(h)** Comparative craniofacial effects captured by Neandertal-aligned and ancestral-aligned shape axes, highlighting a composite of both axes. **(i)** Validation of Neandertal-informed shape effects using canonical variate analysis on a sparse craniofacial landmark dataset from hominin skulls. Shapes from (h) are projected into this space, with corresponding axes indicated.

To assess whether the genetic effects on the chin and midface are coupled, we recomputed the ancestral-derived phenotypic axis using only SNPs associated with midface or chin morphology (r^2^ < 0.1; *P* < 5e-8; facial segments 4 and 11 respectively). Derived alleles (*n* = 1154) at loci associated with midfacial shape also increased chin prominence (**Fig 3c** right), and conversely, derived alleles (*n* = 247) at loci associated with chin shape reduced midfacial projection (**Fig 3c** left). Thus, in both directions, the derived state tended to produce a more prominent chin together with a flatter, less prognathic midface. These bidirectional effects indicate that the genetic architecture underlying these traits is closely aligned.

### Introgressed variants recapitulate Neandertal skull features in modern humans

Next, we investigated how alleles introgressed from Neandertals ∼50 *kya* contribute to craniofacial diversity today. A list of derived alleles with high confidence of being introgressed was obtained from Wei *et al.*^25^ (**Fig 3a**) and filtered down to 79 quasi-independent (r^2^ < 0.1) SNPs that reached *P* < 5e-8 in our craniofacial GWAS (whole craniofacial surface). Using our context-aware approach in the UKBB sample (*n* = 50,622), we obtained a Neandertal-aligned axis from the homozygous introgressed states, capturing 4.65% of shape variance. Facial features associated with an increase in introgressed alleles were visualized at the distribution margins of our UKBB sample (**Fig 3d**, Fig S3a). The relative confidence of local features was assessed using vertex-wise R^2^ (**Fig 3e**), highlighting a higher confidence in the upper and midface, including the nose, and aspects of the cranial vault (maximum R^2^ = 0.60%). The lowest confidence was observed at the lower face, notably the chin (minimum R^2^ = 0.20%). Many of the associated craniofacial features match fossil evidence of Neandertal skulls^43^, including a prominent supraorbital torus, receding frontal bone, a wider nose, more prognathic midface, and flatter zygomatic arches (**Fig 3f**). Furthermore, the cranial vault was elongated with the point of greatest breadth positioned more posteriorly, in contrast to the more globular vault seen on the UKBB mean shape (**Fig 3g**). A clustering of individual SNP effects suggests that different features may be influenced by distinct sets of introgressed alleles (Fig S3b). These results shed light on the lasting effects of Neandertal introgression on the craniofacial traits of present-day humans.

Modelling the directional effects of Neandertal introgressed alleles on the genetic background of modern humans yielded a virtual hybrid of both, showing more modern features in regions with a lower predicted contribution. To address this, we modelled the effects of introgressed SNPs on a genetic background with a higher proportion of ancestral alleles, then compared the resulting morphologies to a revised landmark dataset of hominin skulls, including modern *Homo sapiens* and Neandertals^44^ (Fig S3c). We extended our analysis of the face above to the full craniofacial surface by first recomputing the ancestral-aligned shape axis using 2869 quasi-independent SNPs at craniofacial GWAS loci (r² < 0.1; *P* < 5e-8). The axis captured 6.88% of shape variance. Notably, the ancestral-aligned and Neandertal-aligned axes were positioned at a 90.1° angle, indicating that they captured largely uncorrelated features in our UKBB sample (**Fig 3h**). The composite gestalt (SAN) showed increased prognathism and a reduced chin, traits consistent with Neandertal facial morphology reported in the literature^43^.

To further validate that these axes reflected known morphological differences, we analysed cranial landmark data from 245 modern skulls (21 populations), 8 Neandertals, and 15 other extinct *Homo*. We sampled 100 landmarks spanning the maxilla, orbits, and neurocranium (**Fig 3i**), focusing on regions of minimal soft-tissue thickness, and indicated homologous landmarks on the modelled craniofacial surfaces (Supplementary Note and Fig S3c). Canonical variate analysis revealed that the first variate (CV1) captured differences between *Homo sapiens* and other *Homo* (**Fig 3i**, Fig S3d), while the second (CV2) reflected separation of Neandertals from early *Homo*. Projecting the genetically constructed gestalts from **Fig 3h** into this space, along with their associated axes, revealed strong alignment with skull-based shape differences between *Homo sapiens* and Neandertals (21.0° and 3.42° for the Neandertal and composite axes respectively). In a PCA space of the same cranial landmarks, the Neandertal-aligned and composite axes were closer aligned to skull-based shape differences between *Homo sapiens* and Neandertals compared to random axes (*P* = 3.90e-3 and *P* = 9.90e-3 respectively; empirical test; see Supplementary Note). Collectively, these findings indicate that Neandertal introgression has left a lasting craniofacial imprint in present-day humans, and that the effects modelled using our framework align with known skeletal differences between *Homo sapiens* and Neandertal skulls.

### Genetic liability-aware modelling reveals endophenotypes of clinical conditions

Risk variants for etiologically complex clinical conditions often influence cellular dynamics in subtle, quantitative ways that are poorly captured by binary case-control outcomes. Endophenotypes are heritable, quantitative traits that exist between genetic predisposition and clinical manifestation and are closely aligned with the genetic effects on developmental mechanisms, whether the condition has manifested or not. We sought to leverage genetic risk variants identified in independent GWAS datasets^22,31,45^ that overlapped with craniofacial shape loci to model craniofacial endophenotypes and demonstrate that they can stratify unseen cases from controls. Our analyses focused on one of the most prevalent congenital anomalies, cleft lip with or without cleft palate (CL/P), which affects 0.1% of all live births^46^, as well as two highly prevalent health conditions: obstructive sleep apnoea (OSA) and myopia, affecting roughly 12% and 23% of the world population, respectively^47,48^.

We identified 48, 807, and 1015 quasi-independent (r^2^ < 0.1) SNPs jointly associated with CL/P and facial shape, BMI-adjusted OSA and facial shape, and myopia and craniofacial shape respectively at a 5% conjunctional FDR (Methods). Next, we used our context-aware framework to define endophenotype axes based on the homozygous states of susceptibility and protective alleles after jointly fitting genetic effect sizes on our UKBB sample (*n* = 50,622).

The susceptibility-associated traits captured by the CL/P endophenotype axis included a wider upper and midface, a wider nasal base, a shorter philtrum, and a compressed lower face (**Fig 4a**). To test whether these were aligned with CL/P case-control status, we compared them to an external dataset of syndromic and non-syndromic facial shape data available from the FaceBase^49^ Consortium [*n* = 3772; 66 genetic conditions with craniofacial dysmorphology, plus CL/P cases (*n* = 76) and unrelated controls (*n* = 54)]. After removing the effects of size, sex, age, and age-squared on facial shape, we constructed a PCA space and projected the CL/P endophenotype axis into this space (Fig S4a and Supplementary Note). Along the axis, cases showed moderate separation from controls (**Fig 4a**; Cohen’s *d* = 0.362, 95% CI = [0.011–0.711]; *P* = 2.15e-2, one-tailed *t* test). Furthermore, the endophenotype axis was more closely aligned with the CL/P cases than with any other of the 66 genetic conditions (Fig S4b–c; *P* = 1.49e-2, one-tailed empirical test), indicating that it captures traits specific to CL/P (Fig S4d).

**Fig 4.**
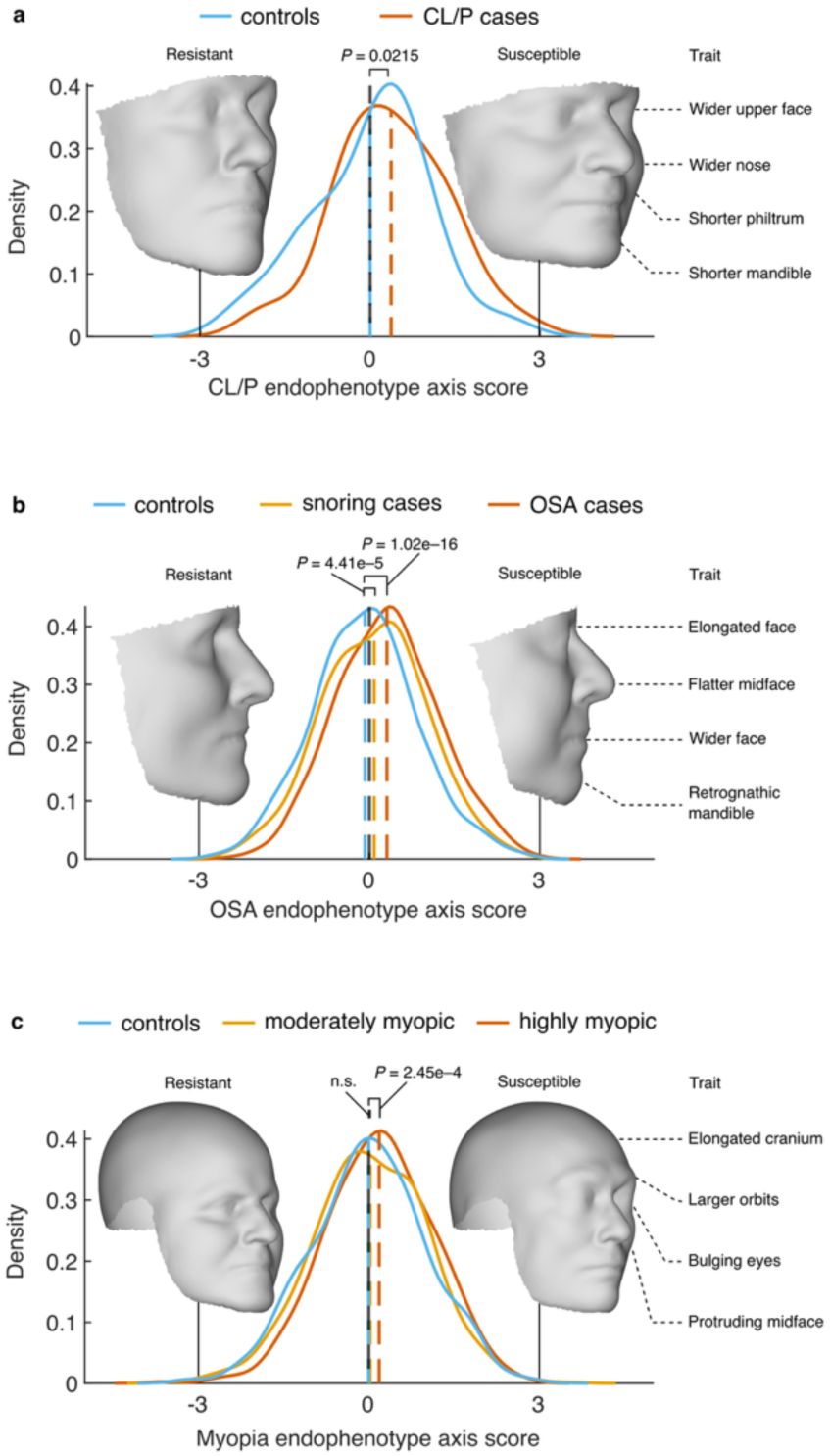
Axes of variation aligned to genetic liability for clinical conditions. Distributions show the one-dimensional projections of new individuals not seen by the endophenotype model. Differences in projection scores between new cases and controls was assessed with a one-tailed *t* test. Visualizations are made at three Euclidean distance units from the UKBB mean shape to illustrate expected variation at the distribution margin. The main susceptibility-associated traits are indicated. **(a)** Cleft lip with or without cleft palate (CL/P)-informed axis indicating distributions of cases (*n* = 76) and controls (*n* = 54). **(b)** Obstructive sleep apnoea (OSA)-informed axis indicating distributions of OSA cases (*n* = 634), snoring cases (*n* = 1000), and non-snoring controls (*n* = 1000). **(c)** Myopia-informed axis indicating distributions of highly myopic cases (*n* = 527), moderately myopic cases (*n* = 1000), and controls (*n* = 1000).

Facial shape is an established risk factor for OSA due to its correlation with upper airway morphology, yet consensus on specific susceptibility traits remains elusive^23^. The OSA endophenotype axis was estimated on a subset of our UKBB sample (*n* = 48,028), leaving out 634 OSA cases along with a random sample of 1000 snoring and 1000 non-snoring controls for validation. The susceptibility phenotype captured by the axis was independent of environmental factors like BMI (Fig S5a) and portrayed a wider, slightly longer, and markedly flatter facial profile with a retrognathic mandible (**Fig 4b**). This facial profile was remarkably consistent with the average facial differences between the unseen OSA cases and non-snoring controls (Fig S5b; *P* < 1e-6, one-tailed empirical test) and both groups could also be stratified based on their endophenotype scores (**Fig 4b**; Cohen’s *d* = 0.422, 95% CI: [0.321–0.522]; *P* = 1.02e-16, one-tailed *t* test). Case-control stratification was much more effective using the whole facial endophenotype, compared to smaller anatomical segments (Fig S5c), suggesting that OSA susceptibility manifests in global facial dimensions. Moreover, facial endophenotype scores showed predictive potential on their own (average ROC-AUC = 0.618) and improved prediction accuracy when age, sex, and BMI were also considered (average ROC-AUC = 0.817; Fig S5d). The controls who reported snoring, a common symptom of OSA, also showed elevated endophenotype scores relative to non-snoring controls (**Fig 4b**; Cohen’s *d* = 0.176, 95% CI: [0.088–0.263]; *P* = 4.41e-5, one-tailed *t* test).

Developmental proximity and structural constraints^50^ suggest a link between craniofacial morphology and myopia, yet the associated craniofacial traits remain poorly defined. A myopia endophenotype axis was estimated on a subset of our UKBB sample (*n* = 48,135), leaving out 527 high and 1000 moderate myopia cases along with a random sample of 1000 controls for validation. The resulting susceptibility phenotype portrayed a protruding upper and midface, a larger vertical orbital opening, bulging eyes, and an elongated cranium (**Fig 4c**). These traits were consistent with the average craniofacial differences between unseen, high myopia cases and controls (Fig S6a; *P* < 1e-6, one-tailed empirical test), and both groups exhibited moderate stratification based on their endophenotype scores (**Fig 4b**; Cohen’s *d* = 0.188, 95% CI: [0.082–0.294]; *P* = 2.45e-4, one-tailed *t* test). Stratification of the high myopia cases from controls was mostly driven by localized morphology around the eyes and orbits (Fig S6b). Notably, the unseen, moderate myopia cases did not stratify from controls (*P* = 0.272, one-tailed *t* test). Altogether, these results indicate significant overlap between genomic loci associated with craniofacial shape and those linked to clinical conditions and demonstrate the construction of genetic liability-aware endophenotypes capable of stratifying unseen cases from controls.

## Discussion

Our study addresses a central question in human biology: how evolutionary, demographic, and health-related genomic signals shape morphological diversity in present-day humans. By analysing craniofacial shape from MR scans of over 50,000 UKBB participants, we derived context-aware axes of shape variation modelled from an expanded catalogue of SNPs associated with craniofacial shape. These axes expose phenotypic configurations that are rare in our White-British UKBB cohort but mirror global patterns of genetic diversity, the evolutionary separation of modern humans from other hominins, the morphological legacy of Neandertal introgression, and risk phenotypes for clinical conditions.

Central to our approach is the view that complex morphological structures embody a high-dimensional, quantitative subset of the phenome, i.e., a continuum of observable traits that can be analysed and decomposed in numerous ways^51^. While linear and angular anthropometric measurements are useful shape descriptors, individual measurements tend to reflect a broad set of influences rather than specific evolutionary or developmental processes^52^. We addressed this by using a bottom-up approach to quantitatively model traits that are aligned with evolutionary and developmental processes, enabling us to meaningfully contextualize human craniofacial diversity. This complements previous studies that have adopted a top-down strategy, leveraging external cues in the phenotypic domain to improve biological alignment and thereby query specific genetic insights. Examples include heritability-enriched traits informed by parent-offspring resemblance^53^, syndrome-aligned traits informed by resemblance to achondroplasia patients^54^, and traits aligned to the shared development of the brain and cranium based on morphological covariation^55^.

Modelling phenotypic axes aligned with allele frequency gradients across global populations revealed continuous trends within our cohort that mirror global patterns of genetic diversity. These findings reinforce that most axes of craniofacial variation, like genetic variation, lie within populations rather than between them, underscoring the fallacy of “racial traits” as biological absolutes and emphasizing a shared evolutionary and developmental basis for human diversity^11^. Interestingly, these axes captured traits consistent with adaptations to local climate, including nasal breadth and alar base features, where temperature and humidity correlate with nares width and internal nasal cavity shape^10^. By incorporating multiple populations from the same and different continents, our results extend earlier demonstrations of continuity between East Asian and European cohorts^13^ and highlight a degree of genetic effect-direction portability across ancestries.

We found that derived alleles at craniofacial loci simultaneously increase chin prominence and reduce midfacial projection, whereas ancestral alleles do the opposite. This genetic coupling provides a parsimonious resolution to a long-standing debate^56^: rather than requiring a specific adaptation for a chin, a mental eminence can emerge as an integrated byproduct of facial retraction^57^. This interpretation aligns with the broader trend of cranial globularization and facial reduction in *Homo sapiens* relative to other hominins, where several craniofacial traits changed together under positive selective pressures^4,28^. These findings position the chin within a covarying developmental field governed by an integrated genetic architecture.

Neandertal fossils reveal the craniofacial differences from *Homo sapiens*^43^, and ancient DNA studies have catalogued introgressed segments in modern genomes^58^. Yet, their aggregate contribution to present-day craniofacial diversity has remained unclear, as several prior studies have focused on single loci^16,24^. We addressed this gap by jointly modelling the directional effects of confidently introgressed variants^25^ associated with craniofacial shape, revealing associations with a broad set of craniofacial traits that closely align with fossil expectations^43^. Notably, Neandertals lack a mental eminence^43^, and introgressed SNPs explained little variation in the chin. Crucially, our aim differed from previous efforts that sought to reconstruct individual Neandertal faces using polygenic models trained on present-day populations^18^. Rather than predicting identities, we model effect directions within a modern genetic background to capture the associated mode of variation, avoiding inconsistencies that arise when projecting polygenic models across large evolutionary distances under divergent gene-environment contexts^59^. Overall, our findings reveal the craniofacial legacy of Neandertal introgression and highlight traits consistent with fossil data.

We identified substantial genomic overlap between craniofacial shape and three prevalent clinical conditions: (i) CL/P, (ii) OSA, and (iii) myopia. These shared genetics were leveraged to derive endophenotype axes aligned with genetic liability that captured recognizable craniofacial configurations and stratified unseen cases from controls. The CL/P axis recovered the broader midface, wider nasal base, and shorter philtrum reported in family-based studies^60,61^. Additionally, the axis indicated a compressed lower face, which challenges previous findings^60^, but aligns with known biological mechanisms such as reduced mesenchyme proliferation in the developing lower facial processes^62^. OSA is a highly prevalent breathing disorder, affecting nearly one billion people worldwide, yet remains critically underdiagnosed and linked to poor health outcomes, underscoring the need for early detection and intervention^48^. Beyond lifestyle factors, BMI, and genetic predisposition, facial morphology is an important risk factor because it covaries with upper airway structure^23^. Our OSA-aligned shape axis captured a facial risk profile independent of BMI, featuring a retrognathic mandible, the most commonly reported facial risk factor, alongside less frequently noted features, such as a flatter, wider, and slightly elongated facial profile^23^. These findings highlight opportunities to improve detection, risk stratification, and targeted interventions such as mandibular advancement devices whose efficacy depends on craniofacial morphology^63^. In the general population, myopia is the leading cause of vision loss worldwide^47^, however, the associated craniofacial traits remain poorly defined. Observations in craniosynostosis, where altered orbital morphology and ophthalmic complications frequently co-occur, indicate a developmental link between craniofacial structure and refractive errors^64^. Here, our myopia-aligned axis emphasized orbital and upper-midface changes consistent with developmental proximity and structural coupling between craniofacial and ocular growth^50^. Interestingly, we identified an elongated, narrowed cranium and upper-midface projection as risk factors for myopia, aligning with the inverse observation of high hypermetropia prevalence in patients with coronal craniosynostosis, who exhibit shorter crania, midface hypoplasia, and shallower orbits^64,65^. Notably, our axis stratified severe but not moderate cases, suggesting that craniofacial morphology may play an underappreciated role in high myopia. Overall, our findings demonstrate that genetic liability for clinical conditions may manifest partially in craniofacial endophenotypes.

While our analyses revealed remarkable continuity in human craniofacial variation, they were based on the genetic and phenotypic diversity present within the UKBB cohort, which consists primarily of older individuals with similar genetic ancestry and self-reported White-British ethnicity. Future studies should therefore include more diverse populations in terms of ancestry and demographics. This is particularly important for clinical applications, where early risk stratification is critical and when risk profiles may depend on population-specific alleles that were not captured in our work because they are absent or non-polymorphic in this sample.

We emphasize that our approach and analyses focus on average phenotypic trends within the cohort, aiming to contextualize morphological diversity in this specific sample. This perspective is fundamentally different from identity prediction, where individuality is embedded in deviations from average trends. We further emphasize that the craniofacial configurations presented in this work illustrate hypothetical extremes, representing expected average morphology at the margins of the cohort’s multivariate shape distribution. While meaningful for highlighting phenotypic trends, they do not correspond to any specific individual, population-level average, Neandertal specimen, or group outside the cohort.

The main input to this work was an expanded catalogue of genetic variants associated with craniofacial shape, initially generated by conducting a multivariate, multiscale GWAS. This initial discovery phase was essential for identifying craniofacial effects beyond the face and establishing a genetic basis that reflects the integrated nature of the craniofacial complex, a critical aspect to the interpretation of evolutionary processes^66^. In doing so, we address a gap left by comparable GWAS focused on facial variation^12,13,18^, which were limited by available imaging and phenotyping methods, and additionally identified roughly ten times as many independent GWS SNPs. While earlier work highlighted a subset of facial shape loci with multiple distinct phenotypic effects^12,18^, our findings instead show that such complexity is ubiquitous, especially near genes with established roles in craniofacial development. Even at individual loci, we reveal additional complexity in spatial regulation by mapping SNP effects to specific anatomical regions and predict diverse *cis*-regulatory strategies through the application of recently developed S2F models^35,36^.

In conclusion, our study demonstrates that the craniofacial shape of contemporary humans embodies genomic signatures of evolution, migration, and health, and that these signals can be disentangled through context-aware axes derived from large-scale imaging and genomic data.

## Methods

### Ethics statement

All participants of the UK Biobank (https://www.ukbiobank.ac.uk/) provided written informed consent. Access to individual-level UKBB data was approved under application number 88320. Local institutional approval (S63179) was granted for this study.

### Image processing and quality control

We obtained 71,220 T1-weighted MR head scans from 66,021 UKBB participants (mean age 64.9 years; 51.9% female) to extract craniofacial shape. Scans were corrected for gradient nonlinearity and intensity bias (N4 algorithm^67^) and denoised via consensus reconstruction from 300 non-linear registrations (*Elastix*^68^) as described in previous work^21^. Head surfaces were isolated by isosurface extraction and mapped to a bilaterally symmetric template comprising 19,734 dense quasi-landmarks using rigid alignment (RANSAC^69^) and non-rigid surface registration (MeshMonk^70^). The regions surrounding the ears and neck were excluded due to their higher complexity and susceptibility to imaging artifacts. Shapes were aligned by Generalized Procrustes Analysis (GPA) and symmetrized.

To identify artefacts, we first curated a sample of 3500 anatomically diverse, artefact-free shapes and used them to determine an empirical per-vertex error distribution to determine whether vertices were outliers in the rest of the sample using a leave-100-out approach. Once applied to the rest of the sample, shapes with less than 50% intact anatomy were omitted from further analysis and for the remaining shapes, outlier vertices were treated as missing data followed by imputation by reconstruction after incomplete data PCA^71^ on the whole sample.

The resulting vertex configurations were adjusted for the following covariates using partial least squares regression (Matlab 2024b: *plsregress*): sex (field: 31); age and age squared (field: 21003); standing height (field: 12144); weight (field: 12143); position in scanner (field: 25756–25759); assessment centre and date (field: 53, 54); whole head size (field: 25000); centroid size of the symmetrized craniofacial mesh; and the ten first genomic PCs. For all covariates except sex and assessment centre, extreme outliers (at >6 s.d.) were removed, and missing values were mean-imputed.

Since positional variability in the craniofacial configurations is superimposition dependent, we performed a hierarchical segmentation of the craniofacial surface. Initially, the face and cranial vault were separated in correspondence with the underlying viscero- and neurocranium. Since the cranial vault is globular and previous work^21^ confirmed that genetic variants predominantly affect its global dimensions, only a single split was performed by approximately following the coronal suture, yielding anterior and posterior vault segments. Furthermore, given the highly modular nature of facial shape, a data-driven segmentation was performed, guided by a combined similarity matrix (RV and Euclidean distance) to ensure anatomical coherence, yielding a facial segmentation that was highly concordant with previous works^12–14^. For each of the resulting 67 segments, we extracted multivariate shape descriptors using PCA and retained PCs that captured 98% of the variance. Further details on image processing can be found in the Supplementary Note.

### Genotyping and quality control

From the UKBB v3 imputed genotypes (field: 22828), available in BGEN v1.2 format, we extracted individuals with available imaging data who had self-reported White-British ethnicity and clustered in a genetic PCA space (field: 22006). Variants on the autosomes and X chromosome were extracted and filtered for an info score greater than 0.3, MAF greater than 1%, genotype failure rate lower than 5%, and in Hardy-Weinberg equilibrium (*P* value > 1e-6) using PLINK^72^ v2.0. Finally, relatives up to the third degree were removed using KING with a kinship coefficient cutoff of 0.0442. Population structure was captured using the first 10 genomic PCs obtained by using PLINK^72^ v2.0 (--pca) on a pruned set of high-quality autosomal SNPs (genotyping rate > 98%, r^2^ < 0.1 across a 500kb window) located outside known regions of long-range LD^73^.

### GWAS

The final dataset for GWAS consisted of 50,662 individuals (24,261 male and 26,401 female) and 8,922,008 variants. Autosomal variants were encoded using the additive model (0/1/2), whereas variants on the X chromosome were encoded additively in females and hemizygously (0/2) in males. Canonical correlation analysis (CCA) was used to test each SNP against the full set of morphological variables in each craniofacial segment separately. A right-tailed *F* test was used to obtain *P* values.

### Definition of lead variants

First, to assign GWS SNPs to non-overlapping genomic regions, an LD-based clumping algorithm was used, equivalent to the default FUMA^74^ (SNP2GENE) algorithm. Briefly, associations across all 67 craniofacial segments were aggregated into a single GWAS by taking the minimal *P*, and all SNPs were clumped with the most significant SNP in LD using two-stage clumping (r^2^ = 0.6, then r^2^ = 0.1). For these SNPs, LD blocks (r^2^ ≥ 0.6) were estimated, and any two SNPs were merged into the same genomic region if the outermost coordinates of their LD blocks were closer than 250 kb. This resulted in 1175 independent, non-overlapping genomic regions. Next, independently significant signals, referred to as “lead SNPs”, were identified through conditional testing within each region. Any additional lead SNP was required to be genome-wide significant (*P* < 5e-8) for at least one craniofacial segment after adjusting that segment for the genotype vectors of all prior lead SNPs from the same region. This resulted in 2579 independently significant SNPs.

### Study-wide significance threshold

To account for the increased multiple testing burden from testing 67 craniofacial phenotypes, a Bonferroni-style study-wide significance threshold was calculated by dividing 5e-8 by the effective number of tests per SNPs. Using 10,000 permutations of 100 randomly selected SNPs, a null distribution of minimal *P* values across the 67 phenotypes was generated. Following the approach by Kanai *et al*^75^, the effective number of independent tests was estimated as 48.2 (s.e.: 2.3), calculated as 0.05 divided by the fifth percentile of the empirical null distribution, resulting in a study-wide significance threshold of 1.04e-9. A total of 2,108 (81.7%) independent lead SNPs were significant at this ⍺ level.

### Replication of GWS SNPs

The replication of GWS variants was performed using independent, previously published GWAS data on facial and cranial vault shape. We selected recent (< 5 years old), independent, large-scale (*n* = 4198–9674) European^12^ and East Asian^13^ whole-face GWAS meta-analyses, and a European whole-cranial vault GWAS^21^. For each lead variant that was missing in any of the replication dataset, the SNP in strongest LD (in-sample; and r^2^ > 0.4) was tested for replication instead. We considered a SNP successfully replicated at a nominal ⍺ level of 0.05. Additionally, a 5% FDR threshold was calculated using the Benjamini-Hochberg procedure.

### LDSC

For each individual shape PC across all 67 craniofacial segments, univariate GWAS summary statistics were generated and LDSC was performed using the HapMap 3 reference list (excluding the human major histocompatibility region) and pre-computed 1000 Genomes Phase 3 European LD scores^33^. The SNP-based heritability of each craniofacial segment was calculated as the sum of its per-PC heritability estimates, weighted by their eigenvalues. Intercepts ranged between 0.98 and 1.04, with a mean of 1.01 (s.d.: 0.01), confirming no residual confounding.

### H3K27ac data processing

H3K27ac ChIP-seq data from a variety of cell types and tissues (*n* = 127) were obtained from public repositories, including ENCODE, SRA, and GEO (Supplementary Data 4). Pre-processing was done following the pipeline described in White *et al*.^12^ Briefly, the raw sequence files (FASTQ format) were aligned to the human genome build 19 using Bowtie2, then sorted and indexed with Samtools. Next, read counts per 10 kb genomic bin were calculated (Bedtools: *coverage*) and normalized to reads per million (RPM). Inter-dataset variability was accounted for by performing quantile normalization (R: *normalize.quantiles*). Genomic bins overlapping with at least one of the 2579 lead SNPs were retained for analysis.

Cell types were grouped into CNCCs (*n* = 12), Carnegie stages (CS13–23; *n* = 22), osteoblasts (*n* = 15), chondrocytes (*n* = 4), and other (*n* = 74). Finally, the 2579 lead SNPs were clustered based on the log-transformed, normalized counts using a weighted k-means++ clustering algorithm (Matlab 2024b: *kmeans*). Weights were proportional to 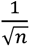 to avoid that the clustering was disproportionately influenced by a specific group. Enrichment of H3K27ac signal in each group relative to the ‘other’ group was assessed by a Mann-Whitney *U* test (two-tailed) on the median signal across SNPs.

### Sequence-to-function predictions

S2F models predicting steady state accessibility in CNCCs or chondrocytes were obtained by running the full ChromBPNet v0.1.1 pipeline with default parameters on a consolidated BAM file of all unperturbed ATAC-seq samples. For CNCCs, we used ATAC-seq data and peak set from Naqvi *et al*^76^. For chondrocytes we used ATAC-seq data from Long *et al.*^77^ and called ATAC-seq peaks using macs2. Five independent train-test-validation splits were used.

S2F models predicting SOX9 or TWIST1 responsiveness (ED50 or full depletion effects) were obtained as described in Naqvi et al^35^. Each of the five steady-state CNCC ChromBPNet models (one from each training split) was fine-tuned with the effect size of full TF depletion or ED50, as defined in Naqvi et al^35^. Full depletion effects (log2 fold-change in ATAC-seq signal) for all 151,457 ATAC-seq peaks was used for fine-tuning, whereas for ED50 fine-tuning was limited to the set of 35,712 SOX9-dependent ATAC-seq peaks or the set of 50,850 TWIST1-dependent ATAC-seq peaks as defined in Naqvi *et al*^35^. Learning rate was set to 1e-3 as in the original ChromBPNet training. Models were trained for 50 epochs but had an early stopping callback with patience of 5 epochs. The best-performing model (lowest loss on the validation set) was used. The same loss functions as the pretrained model were used, except the weight for the multinomial NLL loss (for the base-resolution profiles) was set to 0. Reverse-complemented sequences were used as data augmentation.

SNP effects on any of the seven molecular phenotypes predicted by the seven S2F models were obtained using the ChromBPNet *variant-scorer* function (https://github.com/kundajelab/variant-scorer). Predictions were made for each fold and averaged across all five folds for each model. For selected SNPs at the *OSR1* locus, contribution scores in a 200bp window around each SNP were generated for the reference and alternate alleles using the *variant_shap.py* function.

Predictions were normalized between the S2F models by using a rank-normal transformation and subsequently squaring the data, effectively aligning it to a one degree of freedom χ^2^ distribution. The latter was done as the sign of the prediction is tied to the arbitrary choice of reference allele and to produce distributions with heavier tails as in the raw predictions. Hierarchical clustering was performed on GWS SNPs with substantial predictions (*P* < 0.01) in or near the combined set of CNCC and chondrocyte ATAC peaks (≤ 750 bp) based on pairwise Euclidean distance and Ward linkage (Matlab 2024b: *linkage*).

### Ancestral-derived alleles

Ancestral alleles at each SNP were obtained from the 1000 Genomes^41^ Phase 3 variant call format (VCF) files, where the ancestral state at each SNP was inferred from the six-primate Enredo-Pecan-Ortheus (EPO)^42^ alignments available in Ensembl v71. Only SNPs with high-confidence calls were intersected with GWAS SNPs, meaning that the human ancestral state inferred from the human-chimp-macaque alignment agrees with the human-chimp ancestor and the chimp. After intersecting the resulting list with GWS SNPs for the whole craniofacial surface, whole face, midface (facial segment 4), and chin (facial segment 11), SNPs were pruned using in-sample LD (r^2^ < 0.1) whilst prioritizing the most strongly associated SNP (PLINK^72^ 2.0: *--clump*), yielding 2869, 1958, 1154, and 247 approximately independent predictor SNPs for the respective craniofacial segments.

### Neandertal introgressed alleles

A list of 235,592 Neandertal informative mutations was obtained from Wei *et al*.^25^ who expanded the list from Sankararaman *et al*.^58^ to incorporate strongly linked (r^2^ > 0.99 and within 200 kb) SNPs in the UK Biobank imputed dataset. After filtering for SNPs that reached *P* < 5e-8 in the GWAS, had a MAF > 1%, and whose derived allele was not observed in any of the YRI (*n* = 108) samples from the 1000 Genomes Phase 3 dataset^41^, 2131 SNPs were retained. A set of 79 approximately independent predictor SNPs was obtained by pruning this set using in-sample LD (r^2^ < 0.1) whilst prioritizing the most strongly associated SNP in the GWAS (PLINK^72^ 2.0: *--clump*).

### Population informative SNPs

All GWAS SNPs were tested for differences in allele frequency between the GBR and each of the other populations in the 1000 Genomes Phase 3 dataset^41^ using Fisher’s exact test (right tailed χ^2^). We chose GBR since they were genetically closely aligned with our UKBB sample (Supplementary Data 5). The resulting *P* values were consistent with Wright’s FST values, calculated using PLINK^72^ 2.0 (*--fst*). Given the different degrees of differentiation between intra- and inter-continental populations, sets of approximately independent predictor SNPs were selected slightly differently in both cases.

For non-European populations, SNPs were first filtered to have *P* < 5e-8 in both the GWAS on the whole craniofacial surface and in the scan for allele frequency differences, whereas for European populations, conjunctional FDR analysis^78^ was applied to jointly reject both null hypotheses instead, retaining SNPs at a 5% threshold. Utah residents with Northern and Western European ancestry (CEU) yielded no differentiated SNPs relative to GBR and were excluded from downstream analysis. To obtain quasi-independent predictor SNPs, the most strongly associated GWAS or conjunctional FDR signal was prioritized when pruning for LD as above.

### Susceptibility alleles

GWAS meta-analysis summary statistics of cleft lip with or without cleft palate (*n* = 170 cases, 988 case-parent trios, and 835 controls) were obtained from Leslie *et al*^22^. BMI-adjusted GWAS meta-analysis summary statistics of binary sleep apnoea status in FinnGen and the Million Veterans Project (MVP; *n* = 749,890) were obtained from Kurniansyah *et al*^31^. GWAS summary statistics for binary myopia status in the MVP (*n* = 65,574 cases and 333,242 controls) were obtained from Verma *et al*^45^. These studies were carefully selected to avoid overlap with our UKBB sample. Conjunctional FDR analysis^78^ was applied to jointly test if a SNP was associated with craniofacial (whole craniofacial surface) or facial shape (whole face) and the condition. Based on prior knowledge from the literature, we focused on the face for CL/P and OSA and examined the broader craniofacial surface for myopia. To obtain quasi-independent predictor SNPs, the most strongly associated conjunctional FDR signal was prioritized when pruning for LD as above.

### Context-informed shape axes

Case-control shape axes capturing the craniofacial effects associated with case-control status were obtained by connecting the average shapes of cases 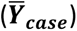 and controls 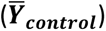. The axis is represented by its unit vector, denoted 𝒖_𝑪𝑪_(***Eq. 1***).

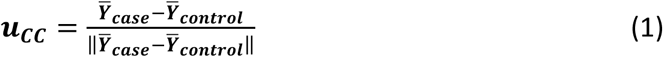

To estimate a polygenic shape axis, a set of 𝑁 SNPs (𝑥_!_, 𝑥_2_, …, 𝑥_3_) was regressed onto the segment-specific PC scores (𝒀) in the full GWAS sample (*n* = 50,662) using partial least squares regression (Matlab 2024b: *plsregress*) under the additive genetic framework (***Eq. 2***). Consistent with the GWAS, we retained PCs explaining 98% of shape variance. No additional covariates were included, as all shape data had been pre-adjusted prior to PCA. Only SNPs on the autosomes were included in the model.

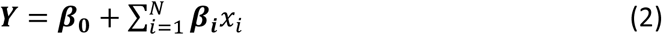

Polygenic shape axes were obtained slightly differently depending on the application. In the case of population-differentiated SNPs, fitted shapes for population 𝑃, denoted 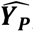 were obtained by evaluating the model at twice the population mean allele frequency for each SNP, denoted 𝑎_6,:_ (***Eq. 3***).

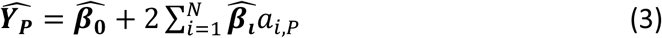

The corresponding phenotypic axis was obtained by connecting 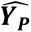 with the fitted values of our UKBB sample, which is simply the mean shape 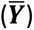. The unit vector along this axis is denoted 𝒖_𝑷_(***Eq. 4***).

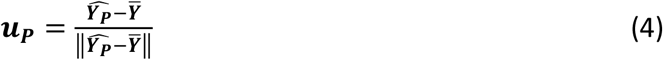

To construct polygenic shape axes for other purposes, including Neandertal introgressed, ancestral, derived, and risk alleles, SNPs were first recoded such that 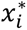 reflected the count of the respective allele type (***Eq. 5***).

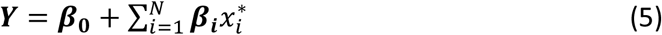

The model was subsequently evaluated at the homozygous states (e.g., two ancestral versus zero ancestral alleles at each SNP), yielding fitted values 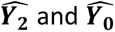 used to define the shape axis with its unit vector 𝒖_𝑯_ (***Eq. 6***).

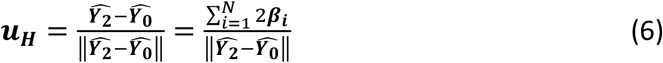

For all purposes, differential gestalts used for visualizing affected anatomy were displaced by 𝑘 Euclidean distance units along the phenotypic axis (***Eq. 7***) with values for 𝑘 indicated throughout the results. We opted to use unit displacements to more straightforwardly produce phenotypes with comparable deviations from the mean.

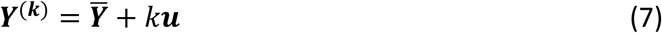

To obtain an individual-level score, denoted 𝑠_6_ for individual 𝑖 along any shape axis, the individual’s shape, denoted 𝒚_𝒊_was projected onto the axis via the dot product with its unit vector (***Eq. 8***).

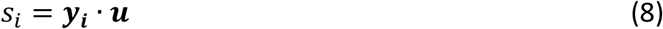

### Case-control status

Within our UKBB GWAS sample, sleep apnoea cases (*n* = 634) were identified using the ICD-10 code G47.3 (field: 41270 — “*Sleep apnoea*”). Snoring cases (*n* = 17,146) were obtained from the UKBB touchpad questionnaire data (field: 1210 — Q: “*Does your partner or a close relative or friend complain about your snoring?*”). Individuals without the ICD-10 code for sleep apnoea and who reported no snoring were considered controls (*n* = 30,665). Additionally, individuals were grouped as highly myopic (dioptre ≤ –6.00; *n* = 527), moderately myopic (–0.75 < dioptre ≤ –6.00; *n* = 3114), or control (*n* = 7110) as in Guggenheim *et al*^79^ (field: 20262). Logistic regression (matlab 2024b: *fitglm*) of the shape PCs onto case-control status, followed by a deviance test (right tailed χ^2^) was used to test the validity of (cranio)facial shape as a biomarker for case-control status.

### 1KG ancestry space

SNPs were intersected across UKBB and the 1KG dataset (*n* = 2504 individuals) and were retained if they had a MAF > 1% and a genotyping rate > 98% in both datasets. A genetic ancestry space was constructed with PLINK^72^ v2.0 (*--pca*) using a pruned set of autosomal SNPs (r^2^ < 0.1 across a 500kb window) located outside known regions of long-range LD^73^. A randomly selected sample of 2500 UKBB participants was projected into the PCA space.

### Phenotypic subspace of axes aligned to population divergence

For each of the 24 1KG populations (excluding GBR and CEU), we constructed a corresponding phenotypic axis and sampled the expected shapes at three positive Euclidean distance units along that axis. Because all sampled shapes exhibited similar magnitudes of deviation from the UKBB mean shape, we rescaled these differences to reflect population divergence by multiplying them by the population-level FST between GBR and each respective 1KG population. Finally, we generated the phenotypic subspace by performing PCA on the recalibrated data.

### Global and local normal displacements

Global and local shape differences between two meshes were quantified by computing displacements along the surface normals at each vertex. Unless stated otherwise, the UKBB mean shape was used as the reference. For global differences, normal displacements were calculated once after a Procrustes superimposition using all vertices. For local differences, neighbourhoods cantered on each vertex and spanning 10% of the craniofacial mesh were defined based on mesh adjacency. Each corresponding neighbourhood pair from the two meshes was then Procrustes superimposed, and normal displacements were calculated at the centre vertex.

## Data availability

UK Biobank data is available for *bona fide* researchers worldwide for health-related research in the public interest, accessed through a formal application process via the UK Biobank website (https://www.ukbiobank.ac.uk/). The FaceBase (https://www.facebase.org/) data is available to researchers via controlled access under accession number FB00000861. GWAS summary statistics for all 67 craniofacial segments are available from GWAS Catalog (https://www.ebi.ac.uk/gwas/home) upon publication of this work. The CL/P, OSA, and myopia GWAS summary statistics are available from GWAS Catalog under accession numbers GCST90652505, GCST90693190, and GCST90475880 respectively. An overview of publicly available H3K27ac datasets is available in Supplementary Data 4. The 1KG dataset is freely available at (https://ftp.1000genomes.ebi.ac.uk/vol1/ftp/release/20130502/). A list of Neandertal-introgressed variants is available from Wei *et al*.^25^ on Github (https://github.com/AprilWei001/NIM/blob/main/Other/expandedNIM.tags). Raw sequence files for training S2F models are available from Gene Expression Omnibus (GEO: GSE267008, GSE205904). Processed sequence data and pre-trained S2F models are available from Naqvi *et al*.^35^ on Zenodo (https://zenodo.org/records/14633030).

## Code availability

KU Leuven provides the MeshMonk v.0.0.6 spatially dense facial-mapping software, free to use for academic purposes available at (https://github.com/TheWebMonks/meshmonk). The latest version is available from the FigShare repository of a previous publication (https://doi.org/10.6084/m9.figshare.c.6858271.v1). Matlab implementations of the hierarchical spectral clustering to obtain facial segmentations are available from a previous publication (https://doi.org/10.6084/m9.figshare.7649024.v1). The following software packages are available from Github: conditional FDR (https://github.com/precimed/pleiofdr), ChromBPNet (https://github.com/kundajelab/chrombpnet), variant-scorer (https://github.com/kundajelab/variant-scorer). The statistical analyses in this work were based on functions in Matlab 2024b, Python 3.12, R v4.5.1, PLINK 2.0, MeshMonk v0.0.6, GREAT v4.0.4, LDSC v1.0.1, ChromBPNet v0.1.1, R packages (geomorph 4.0.10, MASS 7.3-65, Morpho 2.13, ggplot2 3.5.2, caret 7.0-1), and Python packages (SimpleITK 2.5.3, Open3D 0.19.0, NumPy 2.3.4, SciPy 1.15.3, joblib 1.5.2).

## Acknowledgements

We are grateful to the participants, medical staff, and researchers of the UK Biobank who made this work possible. This research has been conducted using the UK Biobank Resource under application number 88320. Part of the resources and services used in this work were provided by the VSC (Flemish Supercomputer Center), funded by the Research Foundation - Flanders (FWO) and the Flemish Government. This project was supported in part by the National Institute of Dental and Craniofacial Research (R01DE027023), Research Fund KU Leuven (BOF GSF/25/038), the Research Foundation-Flanders (FWO G017225N; MSCA SoE FWO 12AD325N), and the Canadian Institutes of Health Research (CIHR 527136).

## Author contributions statement

*Conceptualization:* S. Goovaerts, P. Claes

*Methodology:* S. Goovaerts, J. Devine, N. Claessens, S. Naqvi

*Formal analysis:* S. Goovaerts, J. Devine, N. Claessens, S. Gabbita, S. Naqvi, N. Herrick, K. Pauwels, J. Mohammed, J. Deprest, T. Kivisild

*Investigation:* S. Goovaerts, J. Devine, N. Claessens, P. Claes, S. Naqvi, S. Gabitta, K. Pauwels, T. Kivisild, J. Mohammed

*Data Curation*: S. Goovaerts, J. Devine, N. Claessens, J. Deprest, S. Gabitta, T. Sofer, B. Hallgrímsson, A. Mounier, N. Herrick, S. Bauermeister

*Visualization:* S. Goovaerts, J. Devine, S. Naqvi, S. Gabitta

*Supervision:* P. Claes, S.M. Weinberg, J. Wysocka, T. Kivisild, S. Naqvi

*Funding acquisition:* P. Claes, S.M. Weinberg, J. Wysocka, M.D. Shriver, J.R. Shaffer, S. Walsh, S.J. Lewis, S. Bauermeister, U. Bartsch, S. Richmond, H.K. Long

*Writing - Original Draft:* S. Goovaerts, J. Devine, N. Claessens, P. Claes

*Writing - Review & Editing:* All authors

