## Supplementary Information for "The craniofacial shape of modern humans embodies genomic signatures of evolution, diversity, and clinical conditions"

**This PDF includes:**

Supplementary Note
Fig S1–6

**Other Supplementary Files for this manuscript include the following:**

Supplementary Data 1–5

### Supplementary Figures

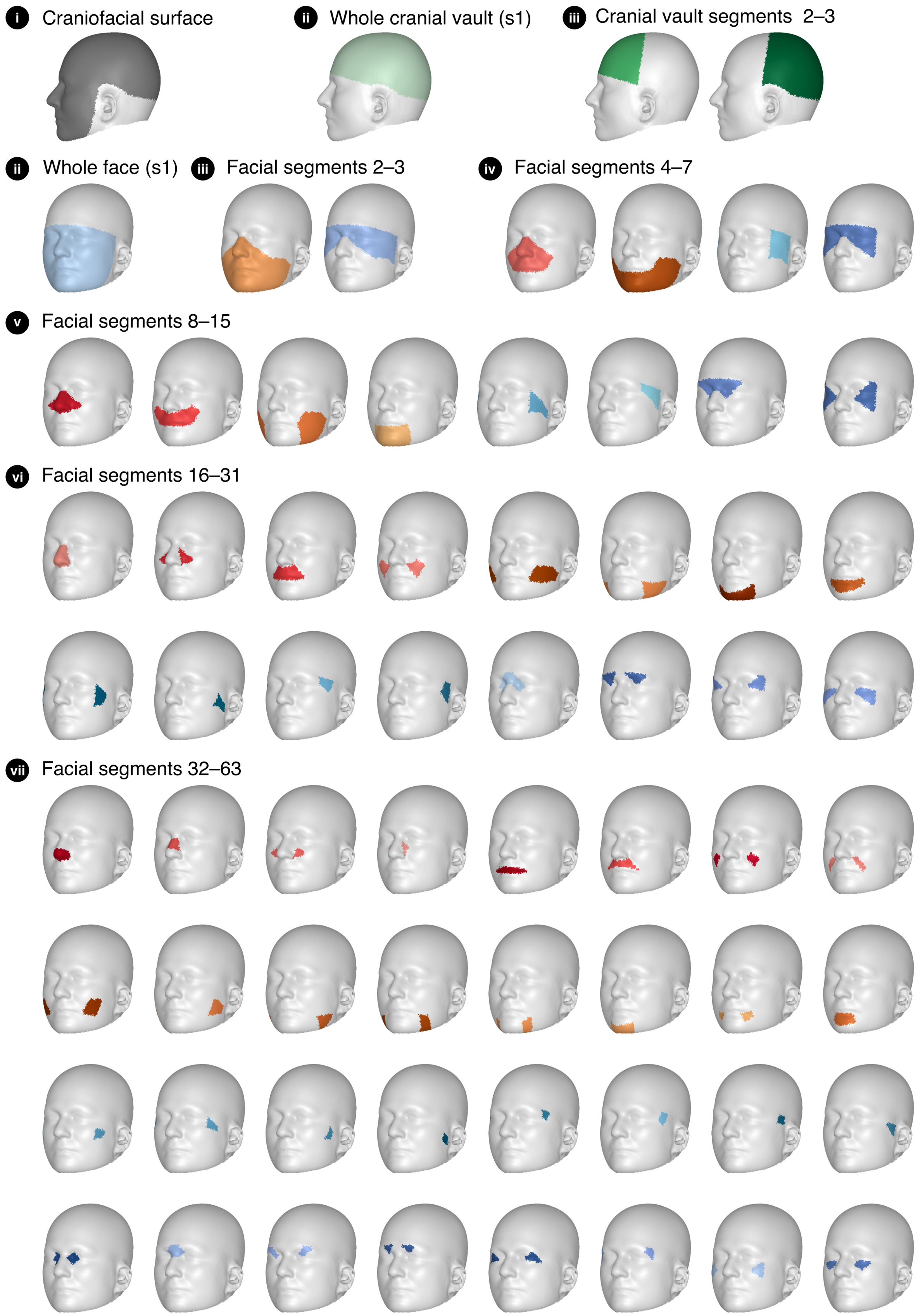


**Fig S1 Overview of craniofacial segments.** Roman numerals indicate the hierarchical segmentation level, with "i" indicating the whole craniofacial surface. Colours match Fig 1.

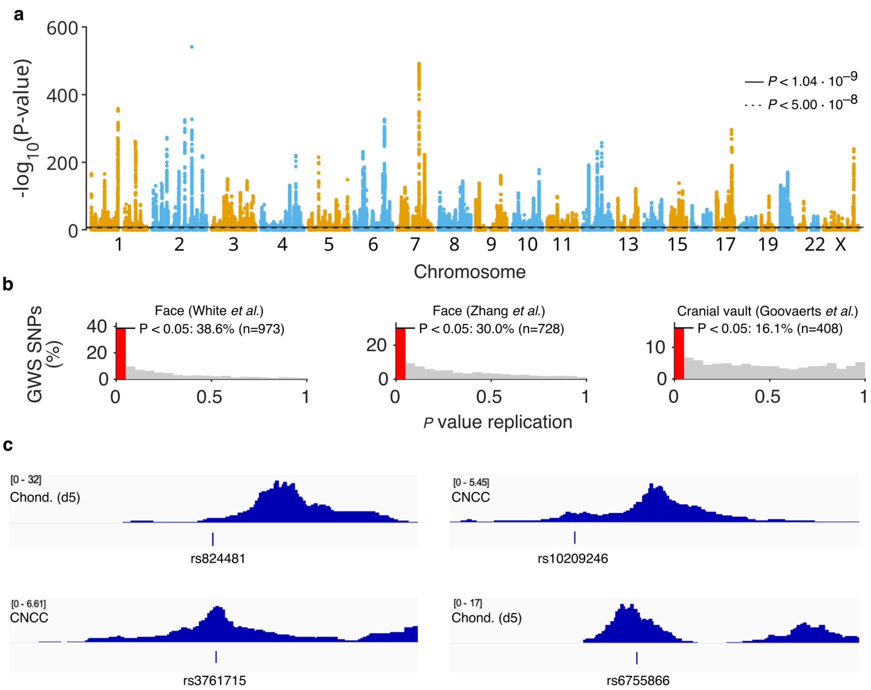


**Fig S2 Additional information on GWAS and replication. (a)** Manhattan plot of craniofacial GWAS, depicting the minimum *P* value across the 67 segments. **(b)** Replication *P* values in three previous GWAS datasets. The red bin ranges from [0, 0.05] and hence depicts the percentage of SNPs replicating at $\alpha = 0.05$. **(c)** ATAC signal in CNCCs or chondrocytes at day 5 (d5) near 4 SNPs at the *OSR1* locus for which we performed functional predictions in Fig 1.

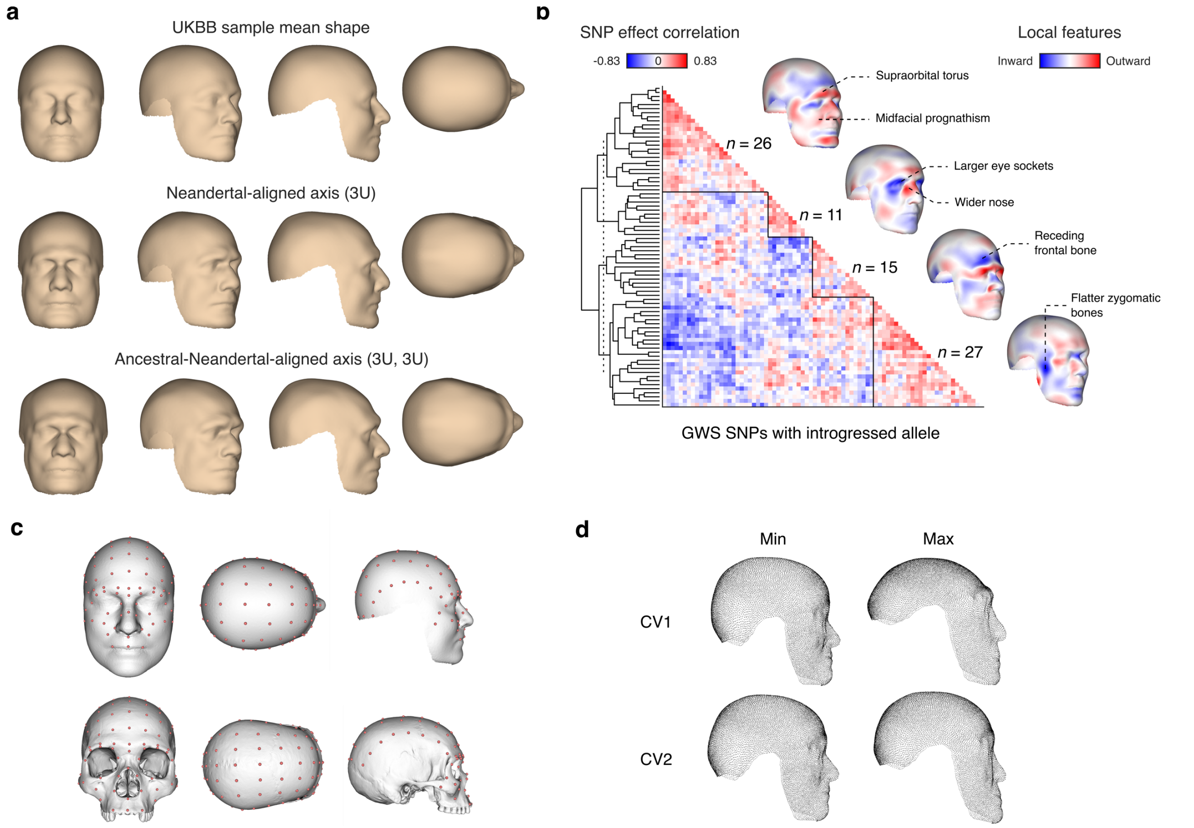


**Fig S3 Additional information on Neandertal-aligned shape axis. (a)** Comparison the UK Biobank (UKBB) mean shape and the expected shape at the UKBB distribution margins (±3 Euclidean distance units) along the Neandertal-aligned shape axis modelled from introgressed SNPs (*n* = 79). **(b)** Hierarchical clustering of introgressed SNP effects (*n* = 79). Clustering was performed on the pairwise correlation matrix of SNP coefficients. Craniofacial heatmaps depict local normal displacements relative to the UKBB mean shape. **(c)** Overview of sparse landmarks used for comparative analysis of surface and skull morphology. **(d)** Effect on surface morphology associated with the canonical variate 1 and 2 in Fig3i. Dense-landmark configurations were obtained by performing a thin plate spline interpolation on the sparse landmarks shown in (c).

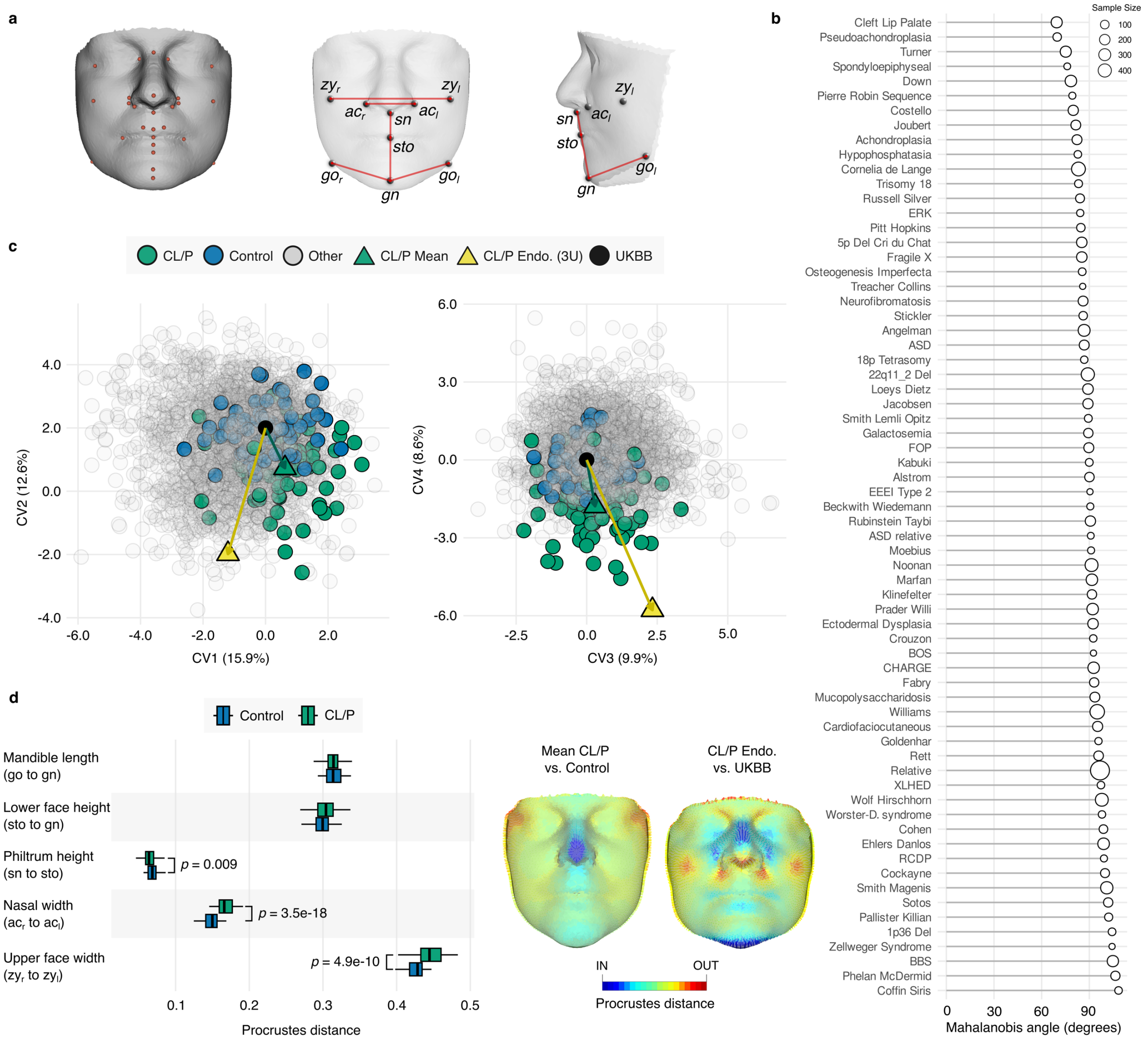


**Fig S4 Validation of CL/P endophenotype axis. (a)** Schematic of sparse landmarks used to project the UK Biobank (UKBB) mean and CL/P endophenotype axis into the shape space of FaceBase (left). Clinically relevant linear distances are also shown (*gn*, gnathion; *go*, gonion; *zy*, zygion; *sn*, subnasale; *sto*, stomion; *ac*, nasal alar crest) (middle, right). **(b)** Mahalanobis angle between the CL/P endophenotype axis and the mean shape of every genetic condition in FaceBase with >10 individuals (n = 3772), sorted from lowest or most similar (top) to highest (bottom). Sample size is denoted by the circle size. **(c)** Canonical Variate (CV) analysis of principal component scores across 67 genetic conditions ($n$ = 3030). The UKBB-CL/P endophenotype axis (yellow) and UKBB-CL/P mean axis (green) are superimposed on distributions of CL/P patients (green), unaffected unrelated controls (blue), and all other genetic conditions (grey). **(d)** Linear distances between key landmarks for controls (blue) and CL/P patients (green) are shown (left) alongside mean shape differences between the CL/P case-control axis, as well as between the CL/P endophenotype-UKBB template axis (right).

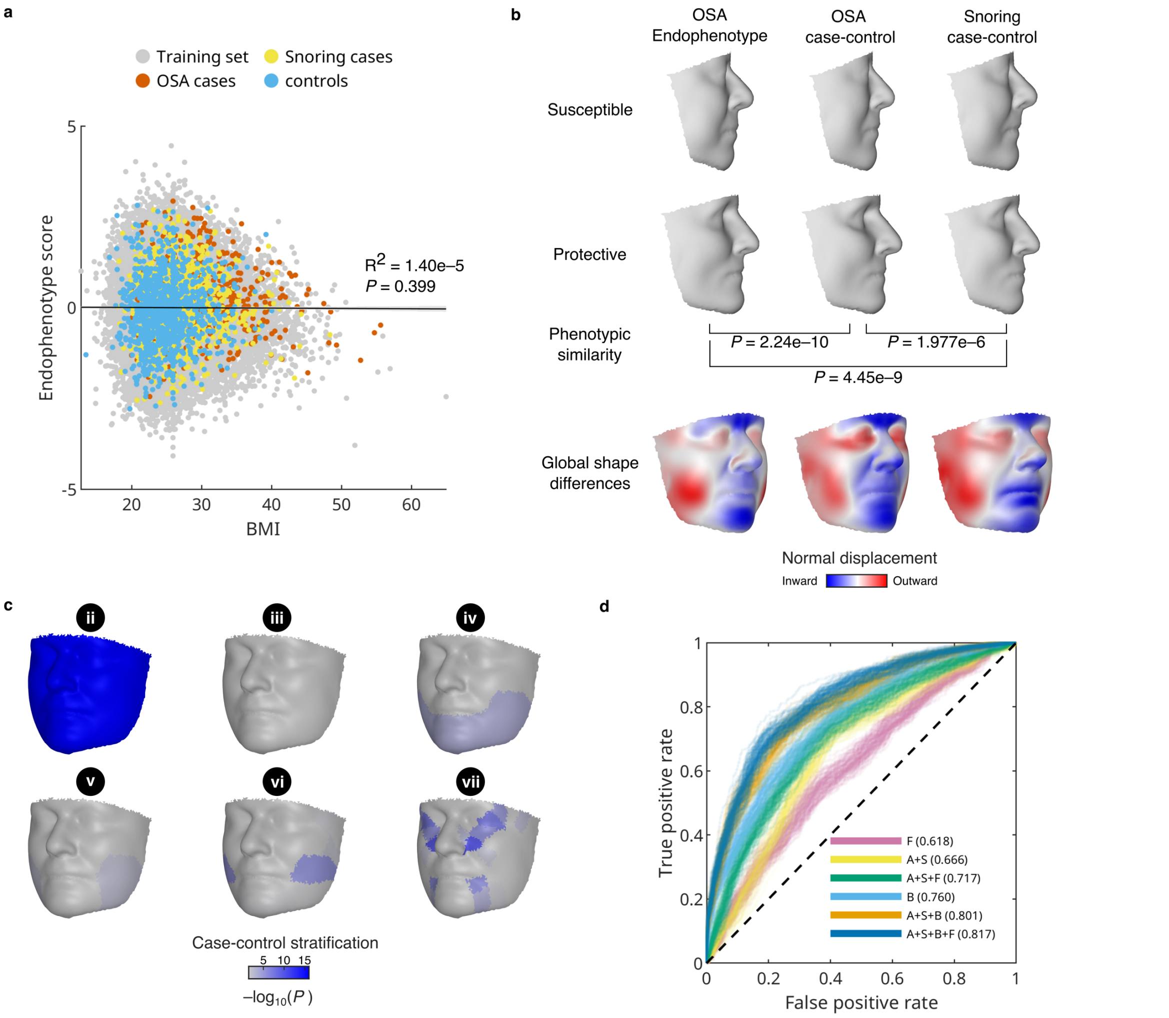


**Fig S5 Validation of OSA endophenotype axis. (a)** Facial endophenotype scores are not correlated with BMI (*n* = 50,622; *P* = 0.399; *F* test). **(b)** Similarity of the obstructive sleep apnoea (OSA) endophenotype with OSA case-control and snoring case-control differences. All pairwise similarities were significant (one-tailed empirical test on vector angle). Susceptibility and protective gestalts are shown along the associated shape axes at ±3 Euclidean distance units from the UK Biobank (UKBB) mean shape. Global normal displacement heatmaps show shape differences with the UKBB mean shape. **(c)** *P* values for case-control stratification when using different facial regions (two sample *t* test; *n* = 634 cases and *n* = 1000 controls). Roman numerals indicate the hierarchical segmentation level, 'ii' indicates the whole face. **(d)** ROC curves for OSA prediction models using logistic regression (For details see Supplementary Note; F = facial endophenotype; A = age; S = sex; B = BMI). Average area under the curve for 100 random train/test splits (50–50%) is indicated for each model.

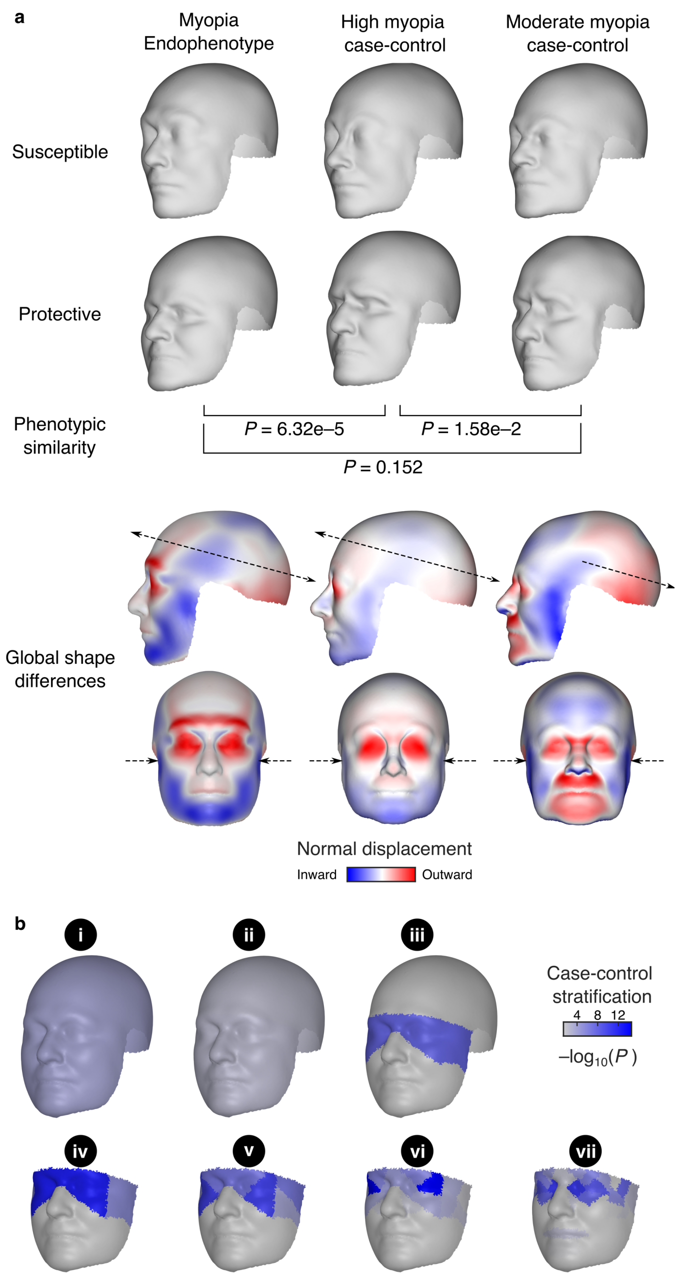


**Fig S6 Validation of myopia endophenotype. (a)** Similarity of the myopia endophenotype with high myopia case-control and moderate myopia case-control differences. All pairwise similarities were significant (one-tailed empirical test on vector angle). Susceptibility and protective gestalts are shown along the associated shape axes at ±3 Euclidean distance units from the UK Biobank (UKBB) mean shape. Global normal displacement heatmaps show shape differences with the UKBB mean shape. Facial narrowing and cranial elongation are highlighted with arrows. **(b)** *P* values for case-control stratification when using different craniofacial regions (two sample *t* test; $n = 527$ highly myopic cases and $n = 1000$ controls). Roman numerals indicate the hierarchical segmentation level, 'i' indicates the whole craniofacial surface.

**Supplementary Note**

*Image preprocessing*

The UK Biobank (UKBB) contains genomic, anthropometric, and health data from ~500,000 British volunteers, and imaging data for a subset, all recruited with informed consent. Under data release v10.1 we obtained 71,220 T1-weighted structural magnetic resonance (MR) scans of the whole head corresponding to 66,021 unique individuals. The dataset was 51.9% female and 48.1% male, with a mean age of 64.9 (s.d.: 7.7), a mean weight of 75.4 kg (s.d.: 15.2), and a mean height of 170 cm (s.d.: 9.4). In total, 84.2% were of self-reported European descent. We subjected this entire dataset to a phenotyping pipeline to extract high-quality craniofacial shape data.

To correct each MR scan for geometric distortions and intensity inhomogeneities, we performed a gradient nonlinearity correction and bias field correction with the N4 algorithm[1]. To minimize noise and imaging artifacts, we implemented an approach from previous work[2], which involved non-linearly registering each volume to 300 other volumes using *Elastix*[3] (*SimpleITK* in Python) and computing a consensus volume based on the median intensity value per voxel. We conducted the registrations with the *Param0000* parameter map (affine and B-spline) using weight, height, sex, and ancestry-matched images. We then extracted the entire head by calculating an isosurface based on voxel intensities and only retained the largest connected mesh to filter out internal structures.

Next, we aimed to establish spatial correspondences to a whole-head mesh template across the extracted head surfaces. This head template, derived from a previously defined mesh[2], was resymmetrized by reflecting the mesh, registering the mirrored version to the original, and averaging the two. The left half was then mirrored across the midsagittal plane, and both halves were stitched along a refined midline, resulting in a symmetric mesh with 19,734 3D quasi-landmarks. We first automatically initialized each surface into the head mesh template space using the Random Sample Consensus (RANSAC) algorithm (*Open3D*[4] in Python). Following these rigid alignments, we non-linearly registered the template to each mesh with MeshMonk[5], resulting in 19,734 homologous, spatially dense vertices (i.e., quasi-landmarks). Ears were ignored during registration due to their higher complexity and susceptibility to imaging artifacts. Part of the back of the head was also cropped out afterwards due to compression artifacts from subjects lying down. The resulting craniofacial surfaces were then aligned to a common shape space via Generalized Procrustes Analysis[6] (GPA; *Morpho*[7] in R) and were subsequently symmetrized.

Since the UKBB imaging protocol was optimized for the brain as opposed to the craniofacial surface, we observed a range of global and local artifacts across the shape data. While global artifacts were related to extensive missing head anatomy, local artifacts manifested as cheek compression, soft tissue accumulation, electrode bumps, and/or minor amounts of missing anatomy (i.e., parts of the chin and/or nose). Our strategy was to detect and remove shapes with global artifacts and impute those with local artifacts. This first involved calculating various mesh and shape metrics (e.g., edge lengths, face areas, Euclidean distances, cosine angles) to train a classifier to screen for high-quality shape data for an outlier detection model. After manually labelling scans with ($n$ = 2,231) and without artifacts ($n$ = 1220), we standardized the mesh and shape metrics (mean: 0, s.d.: 1), split the data into train (85%) and test (15%) sets, and evaluated a series of classifier models (*caret*[8] in R). Using the best performing model (Adaptive Boosting and Bagging, accuracy: 0.80), we predicted all possible artifact-free shapes and sampled anatomically diverse individuals ($n$ = 4,294) from this clean dataset. To maximize diversity, we divided the shapes into 500 cosine angle bins, then randomly sampled and manually verified up to 10 shapes from each bin. Finally, this curated sample was used to inform a principal component analysis (PCA)-based outlier detection model.

To construct the outlier detection model, we first computed empirical per vertex error distributions. Using a cross-validation scheme (leave-100-out), we projected left-out curated shapes into the principal component (PC) space derived from the remaining curated sample, reconstructed these craniofacial shapes, and recorded their vertex-wise Euclidean reconstruction errors. By comparing observed vertex distances to this empirical distribution, we obtained $P$ values and converted these to outlier weights ($w$)

(Eq. 2), which were balanced by a constant (λ) (Eq. 1) with an empirically chosen parameter ($\kappa = 3.5$):

$$\lambda = e^{-\kappa^2/2}/\sqrt{2\pi} \quad (1)$$

$$w = p/(p + \lambda) \quad (2)$$

We updated the weights iteratively for each individual. Over a maximum of $n = 100$ iterations, we (1) aligned each shape to the average using a weighted Procrustes superimposition, (2) projected the aligned shape into the curated shape PC space with its current weights, (3) performed reconstruction to vertex coordinates, (4) calculated vertex-wise Euclidean distances to the reconstruction to flag abnormal anatomy, and (5) repeated this process until the weights stabilized ($w < 0.0001$ for any updated vertex) or $n$ iterations were reached. We also incorporated a small regularization coefficient ($10^{-12}$) during the projection step to prevent the model from reconstructing extreme shapes and artifacts[9].

To identify and remove global outliers, we performed PCA on the outlier weights, clustered the variation with K-means ($K$ = 6) (*kmeans* in MATLAB), and removed one cluster ($n$ = 2454) with misaligned surfaces due to extensive missing anatomy. To maximize sample size whilst retaining the same number of individuals across all segments (see *Image segmentation*), we defined vertices with an outlier weight $w < 0.5$ as outliers and treated them as missing data to impute. Next, we implemented an incomplete data PCA[10] workflow, which involved eigenanalysis on the covariance matrix estimated from vertices with known values. The resulting ordination scores were computed as:

$$u_{jk} = \sum_{i=1}^{n} \omega_{ij}(x_{ij} - \bar{x}_i)v_{ik} \Big/ \sqrt{\sum_{i=1}^{n} \omega_{ij} \cdot v_{ik}^2} \quad (3)$$

where $u_{jk}$ is the score of observation $j$ on PC $k$, $x_{ij}$ is the coordinate value for variable $i$ and observation $j$, $\bar{x}_i$ is the mean of coordinate $i$ computed from non-missing data, $v_{ik}$ is the loading (eigenvector entry) of coordinate $i$ on PC $k$, $\omega_{ij}$ is a binary indicator denoting whether coordinate $i$ is observed (1) or missing (0) for observation $j$. Every individual's score on a PC is therefore computed from the subset of available vertices, and the denominator scales the score to account for the missing values. Vertices that were marked as an outlier in more than eighty percent of the images were removed before the incomplete data PCA and imputed after PCA reconstruction using thin-plate splines.

*Image segmentation*

Both global and local genetic effects on craniofacial shape have been described in various studies[2,11,12], which tend to analyse the face and cranium separately. To investigate these genetic effects jointly and better contextualize the anatomy, we defined the entire craniofacial surface as the global segment. Next, we partitioned the craniofacial template into the cranial vault and face in correspondence with the underlying viscerocranium and neurocranium, then divided these segments into more localized regions. While the vault was split into anterior (frontal bone) and posterior (parietal, temporal, and occipital bones) aspects, the face was subjected to hierarchical spectral clustering[12] to obtain local, data-driven segments. Like the weighting strategy of Zhang *et al.*[11], we provided a combined matrix, $M = 0.6 \times RV + 0.4 \times distance\ matrix$, as input to the clustering algorithm because it yielded more cohesive segments than the RV matrix alone.

*Neandertal-aligned axis validation*

To validate the Neandertal-aligned axes and contextualize them in hominin evolutionary history, we first downloaded 3D hominin crania landmark data (780 skeletal landmarks) from Mounier *et al.*[13] encompassing a range of populations: seven early Pleistocene specimens, including *H. habilis*, *H. ergaster*, and *H. georgicus*; eight Neandertals from different regions (early, Near East, South Europe, and West Europe); and 245 extant modern humans from 21 populations. Next, we manually sampled 100 landmarks from the dense configuration and identified homologous points on the UKBB craniofacial template surface using juxtaposed 3D viewers in 3D Slicer[14] (Fig S3c). We then leveraged the correspondences in our craniofacial surface dataset and performed barycentric mapping to transfer the UKBB template landmarks to the Neandertal-aligned shape axes. After establishing homology across the cranial landmark datasets, we superimposed the data into a common shape space via GPA (*Morpho*[7] in R). Symmetrization was included to ensure consistency between the skeletal and surface data. To control for allometry and sexual dimorphism in the skeletal data, we regressed out the effects of size and sex (*geomorph*[15] in R). To minimize surface and skeletal landmark placement bias, we subtracted the mean UKBB-skeletal difference vector from each surface to recenter the data.

We next sought to establish a shape space in which the directional affinity of the Neandertal-aligned axes could be assessed relative to the hominin groups. Following the shape adjustments, we defined the skeletal shape data as training data, conducted a PCA to reduce its dimensionality, and projected the Neandertal-aligned shapes axes (Fig 3h) into this space as test data. This PC space (98% variance, 50 PCs) was subsequently utilized to quantify overall shape similarity between the Neandertal-aligned axes and the actual Neandertal group via Mahalanobis angles. Here, we generated an empirical null distribution by calculating Mahalanobis angles ($n$ = 10,000) between random pairs of modern humans and the mean modern human-Neandertal axis, then we measured statistical significance (i.e., one-sided empirical p-value) by comparing the Neandertal-aligned axes to this distribution. To better visualize directional trends, we performed a Linear Discriminant Analysis (LDA). Specifically, the training data were categorized into three hominin groups—Early *Homo* (*H. habilis, H. ergaster, H. georgicus)*, Neandertals, and modern humans—and further compressed to seven PCs (i.e., minimum group size) before training and evaluating the LDA model (*MASS* in R) on the Neandertal-aligned axes. This yielded discriminant functions equivalent to canonical variates (CVs) that enabled visualizations of discriminant trends among the groups. The Neandertal-aligned axes were plotted in CV space alongside the hominins (Fig 3i).

*Cleft lip and palate endophenotype axis validation*

To validate the cleft lip and palate (CL/P) endophenotype axis, we used FaceBase, an external cohort of facial shape data from genetic conditions with craniofacial dysmorphology ($n$ = 3772; 67 groups, minimum $n$ > 10). Because the dimensionality of these data (7160 landmarks) differed from the UKBB facial template, we adopted a landmarking and analysis strategy akin to the Neandertal section. This involved placing sparse landmarks (27 landmarks; Fig S4a) on the UKBB facial template and an existing facial template already mapped to FaceBase[16], then transferring those points to the CL/P endophenotype axis and every individual in FaceBase, respectively, via barycentric mapping. After establishing homology, we superimposed the data into a common shape space via GPA (*Morpho* in R). Symmetrization was included to ensure consistency between the FaceBase shapes and the CL/P endophenotype axis. We also subtracted the mean UKBB-FaceBase difference vector from the UKBB data to recenter it, then regressed out the effects of size, sex, age, and age-squared to control for allometry, sexual dimorphism, and other age-related changes in the FaceBase cohort (*geomorph* in R).

Using the adjusted shape data, we performed PCA and CVA to assess how closely the CL/P endophenotype axis aligned with discriminant trends between all genetic conditions in FaceBase. This first involved projecting the CL/P endophenotype into the FaceBase PCA space as test data, retaining all PCs (100% variance, 41 PCs) due to landmark sparsity, then calculating Mahalanobis angles between the CL/P endophenotype axis and all mean case-control shape axes in FaceBase (Fig S4b). Statistical significance (i.e., one-sided empirical *P* value) was evaluated by treating all non-CL/P angles as an empirical null distribution and comparing the CL/P endophenotype axis angle to this distribution. We additionally visualized the similarity between the actual CL/P case-control axis and the CL/P endophenotype axis in CV space after removing genetic conditions with fewer individuals than variables ($n > 41$; $n = 3030$), training an LDA model on the PCs (*MASS* in R), and evaluating it on the endophenotype axis (Fig S4c). Next, we computed linear distances between key landmarks in control (unaffected unrelated) and CL/P individuals to highlight features consistent with the endophenotype axis (Fig S4d). Two-sample *t* tests were conducted for each distance measure to assess statistical significance. To visualize dense shape similarities between the actual CL/P case-control axis and the CL/P endophenotype-UKBB control axis, we warped the UKBB facial surface to the control mean, CL/P mean, and CL/P endophenotype via thin-plate spline, then calculated the Procrustes distance at each point for both contrasts (Fig S4d).

*OSA ROC analysis*

From the total set of 50,622 samples, we used 48,716 to estimate the OSA-aligned endophenotype axis and performed ROC analysis using only the remaining samples (*n* = 634 cases and *n* = 1000 controls). Endophenotype scores of this hold-out set were calculated by vector projection onto the endophenotype axis (see Methods). Logistic regression models were then trained on a random 50% subset of cases and controls (i.e., *n* = 317 cases and *n* = 500 controls) to predict OSA status using facial scores, BMI, age, sex, or subsets of these predictors. Model performance was evaluated on the other 50%, and this procedure was repeated across 100 random train-test splits for each predictor set.
